# Presynaptic Nrxn3 is essential for ribbon-synapse assembly in hair cells

**DOI:** 10.1101/2024.02.14.580267

**Authors:** Alma Jukic, Zhengchang Lei, Elizabeth R. Cebul, Katherine Pinter, Natalie Mosqueda, Sandeep David, Basile Tarchini, Katie Kindt

## Abstract

Hair cells of the inner ear rely on specialized ribbon synapses to transmit sensory information to the central nervous system. The molecules required to assemble these synapses are not fully understood. We show that Nrxn3, a presynaptic adhesion molecule, is critical for ribbon-synapse assembly in hair cells. In both mouse and zebrafish models, loss of Nrxn3 results in significantly fewer intact ribbon synapses. In zebrafish we demonstrate that a 60% loss of synapses in *nrxn3* mutants dramatically reduces both presynaptic responses in hair cells and postsynaptic responses in afferent neurons. Despite a reduction in synapse function in this model, we find no deficits in the acoustic startle response, a behavior reliant on these synapses. Overall, this work demonstrates that Nrxn3 is a critical and conserved molecule required to assemble ribbon synapses. Understanding how ribbon synapses assemble is a key step towards generating novel therapies to treat forms of age-related and noise-induced hearing loss that occur due to loss of ribbon synapses.

**Summary Statement:** Hearing and balance depend on specialized ribbon synapses that transmit sensory stimuli between hair cells and afferent neurons. We identify Nrxn3 as a key molecular player in ribbon-synapse assembly in hair cells.

## Introduction

In the nervous system, synapses transmit signals between neurons, and proper synapse assembly is critical for the function of neural circuits. While many players have been identified in synapse assembly in the central nervous system, relatively less is known about this process in hair cells, the sensory receptors of the inner ear. Importantly, both noise-induced and age-related hearing loss in humans can result from auditory synaptopathy, or synapse loss, even in the absence of hair-cell death (Liberman, 2017; Wu et al., 2020; Wu et al., 2021). By expanding our current knowledge of the players involved in synapse assembly in hair cells, we can better understand how to re-form these synapses and treat auditory synaptopathy.

Hair cells function to convert auditory, vestibular and lateral-line stimuli into signals that are sent to the brain. In response to sensory stimuli, apical structures called mechanosensory hair bundles are deflected, opening mechanoelectrical transduction (MET) channels and depolarizing the cell. Hair-cell depolarization opens voltage-gated calcium channels (Ca_V_1.3) at the presynapse, resulting in calcium influx that triggers the release of the neurotransmitter glutamate onto afferent neurons (reviewed in: (Fettiplace, 2017)). In order to properly encode sensory stimuli, hair cells use a specialized ribbon synapse for speed and precision (Khimich et al., 2005; Moser et al., 2006). A ribbon synapse is defined by a presynaptic density called a ribbon, which is composed primarily of the protein Ribeye (a splice variant of CTBP2) (Schmitz et al., 2000) (Fig 1D). Ribbons are thought to be important to recruit and tether synaptic vesicles at the presynaptic active zone.

**Fig 1.**
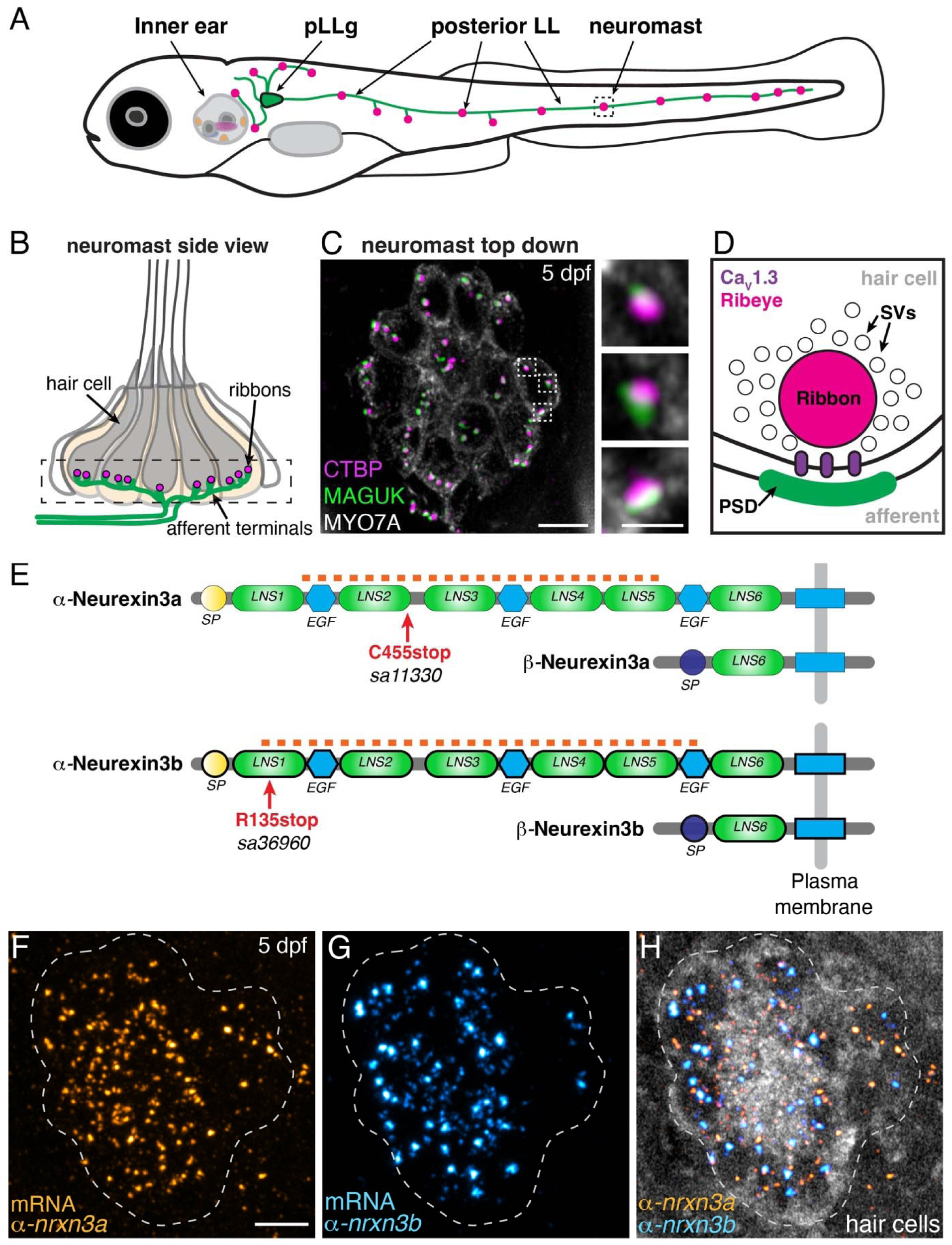
*nrxn3a* and *nrxn3b* are expressed in lateral-line hair cells. **(A)** Schematic showing a larval zebrafish at 5 days post fertilization (5 dpf). Clusters of sensory hair cells are in the inner ear and posterior lateral line (neuromasts, pink). Hair cells of the lateral line are innervated by neurons (green) that project from the posterior lateral-line ganglion (pLLg, blue). (**B**) A lateral-line neuromast organ viewed from the side. Hair cells are labeled in gray, presynapses or ribbons in magenta, and afferent processes beneath the hair cells in green. The dashed box indicates the synaptic layer. (**C**) Immunostaining of the synaptic layer, viewed from the top down. CTBP labels the presynapses or ribbons (magenta), pan-MAGUK labels the postsynapses (green), and MYO7a labels the hair cells (gray). Higher magnification view of three synapses from a single hair cell is shown on the right. (**D**) Schematic highlights the main components of a hair-cell ribbon synapse. The presynapse, or ribbon is composed primarily of Ribeye, a splice variant of CtBP2. The ribbon is surrounded by glutamate-filled synaptic vesicles (SVs). Ca_V_1.3 channels (purple) are clustered beneath the ribbon. AMPA receptors are clustered within the postsynaptic density (PSD). (**E**) There are 2 orthologues of Nrxn3 in zebrafish, Nrxn3a and Nrxn3b. Similar to mammals, each neurexin has a long α form and a shorter β form. We examined zebrafish mutants that are predicted to disrupt the α form of each orthologue (C455stop and R134stop). The α and β forms each have a unique start and signal peptide (SP). Each α form has 6 Laminin G-like domains (LNS) and 3 epidermal growth factor-like domains (EGF). The red dashed line indicates the location of the RNA FISH probes used in F-H. (**F-H**) RNA FISH analysis reveals that both *α-nrxn3a* (F, orange) and *α-nrxn3b* (G, cyan) mRNAs are present in lateral-line hair cells at 5 dpf. In H, hair cells (*myo6b:memGCaMP6s*) are shown in grayscale. RNA FISH labeling was confirmed in 3 independent experiments. The dashed lines in F-H outline the locations of hair cells. All images are from larvae at 5 dpf. Scale bars = 5 µm in C and F, 1 µm in the inset in C.

What molecules are necessary for synapse formation, maintenance and remodeling of this critical sensory circuit remain largely undefined. Cell adhesion molecules (CAMS) are powerful modulators of synapse formation. CAMs act to bridge pre- and post-synaptic domains to initiate, maintain and specify synapses (reviewed in: (Südhof, 2021)). Work in mouse auditory inner hair cells (IHCs) has implicated the CAM Neuroplastin (Np55/Np65) and the neuronal CAM (NrCAM) in the development of IHC ribbon synapses (Carrott et al., 2016; Harley et al., 2018; Newton et al., 2022). More recent work in mouse found that a well-studied family of postsynaptic CAMs, neuroligins (Nlgns), are important for ribbon-synapse assembly (Ramirez et al., 2022). This work found that mice lacking NLGN1/3 have fewer ribbon synapses in IHCs and impaired hearing. Postsynaptic Nlgns classically bind to a family of presynaptic CAMs, called neurexins (Nrxns) (Südhof, 2021). While prior work points to the importance of NLGN1/3 in synapse assembly in hair cells, whether these neuroligins pair with a presynaptic neurexin in the auditory system is not known (Ramirez et al., 2022).

To study the function of neurexins in hair-cell synapse assembly, we used the zebrafish model system. Numerous studies have shown that zebrafish is a relevant model for studying the genetics of hair cells, as many of the core molecules required at hair-cell synapses (ex: Ca_V_1.3 and Ribeye) are conserved between zebrafish and mammals (Brandt et al., 2003; Jean et al., 2018; Lv et al., 2016; Sidi et al., 2004). In zebrafish, hair cells are present in the inner ear and in the lateral-line system (Fig 1A-C). These sensory systems are required for hearing and balance or the detection of local fluid flow, respectively (Gompel et al., 2001; Haddon and Lewis, 1996). Hair cells in the zebrafish inner ear are innervated by neurons in the statoacoustic ganglion (SAG), while hair cells in the lateral-line system are innervated by neurons in the anterior or posterior lateral-line ganglia (aLLg and pLLg) (Zecca et al., 2015).

Our study identifies Nrxn3 as a presynaptic CAM required for hair-cell synapse assembly in both zebrafish and mice. In zebrafish we find that *nrxn3* mutants form ∼60% fewer ribbon synapses in the lateral line and ∼30-45% fewer ribbon synapses in the inner ear. We also show that *Nrxn3* function is conserved in mammals; mice lacking NRXN3 in auditory IHCs form ∼20-25% fewer ribbon synapses compared to controls. In response to stimuli, we find that both pre- and post-synaptic calcium responses in lateral-line hair cells are dramatically reduced in *nrxn3* mutants. Surprisingly, despite fewer synapses and reduced synaptic responses, we observe no detectable deficit in the acoustic startle response, a hair-cell mediated behavior in zebrafish. Overall, our work demonstrates that Nrxn3 plays a crucial and conserved role in ribbon-synapse assembly in mice and zebrafish. This knowledge will inform future research aimed to rebuild synapses and restore hearing after auditory synaptopathy.

## Results

### Loss of Nrxn3 results in fewer ribbon synapses in mature hair cells in zebrafish

Neurexins are classic presynaptic CAMs required for synapse assembly in many contexts (Südhof, 2017). Single cell RNA sequencing (scRNAseq) studies in mice and zebrafish have shown that *nrxn3* mRNA is enriched in hair cells, making it a viable candidate to drive the assembly of ribbon synapses in hair cells (Cai et al., 2015; Elkon et al., 2015; Kolla et al., 2020; Lush et al., 2019; Shi et al., 2023). To determine if Nrxn3 is required for hair-cell synapse assembly, we used the genetically tractable zebrafish model (Fig 1A-D). In zebrafish, there are 2 orthologues of mammalian *Nrxn3*, called *nrxn3a* and *nrxn3b* (Fig 1E). Like mammalian *Nrxn3,* both *nrxn3a* and *nrxn3b* loci are predicted to produce two main isoforms, a long α form, and a shorter β form (Fig 1E) (Gomez et al., 2021). For our analysis we examined the best-characterized α form of *nrxn3a* and *nrxn3b*. We first used RNA FISH, and verified that the α forms of *nrxn3a* and *nrxn3b* mRNA are present in hair cells of the zebrafish lateral line (Fig 1F-H) and inner ear (Fig S1A-B) (Choi et al., 2018).

We next leveraged zebrafish genetics to test whether α-Nrxn3a or α-Nrxn3b is required for the organization of ribbon synapses at 5 days post fertilization (dpf). At this age, the majority of zebrafish hair cells are mature, and both the lateral-line and inner-ear sensory systems are functional (Kimmel et al., 1974; Suli et al., 2012). We obtained existing *nrxn3a* and *nrxn3b* ENU zebrafish mutants from the Zebrafish International Resource Center (Kettleborough et al., 2013). Both alleles have early stop codons that are predicted to specifically disrupt the long α form of each orthologue (Fig 1E). For simplicity, we use Nrxn3a and Nrxn3b from here on to refer to the long α form of each Nrxn3 orthologue, unless otherwise specified.

We first assessed the organization of ribbon synapses in lateral-line hair cells in our *nrxn3* mutants in mature hair cells at 5 dpf. For this assessment, we used immunohistochemistry to visualize presynapses (pan-CTBP) and postsynapses (pan-MAGUK) (Fig 1C). After immunostaining, we quantified the number of complete synapses (paired CTBP-MAGUK puncta), unpaired presynapses (lone CTBP puncta) and unpaired postsynapses (lone MAGUK puncta). Using this approach, we found that there was a slight yet significant reduction in the number of complete ribbon synapses per hair cell in *nrxn3a* or *nrxn3b* single mutants compared to controls (Fig S3A-B; 18.5% reduction in *nrxn3a* mutants and 24% reduction in *nrxn3b* mutants). In *nrxn3b* mutants, but not in *nrxn3a* mutants, we observed significantly more unpaired presynapses compared to controls (Fig S3C). In contrast, we did not observe a difference in the number of unpaired postsynapses in either *nrxn3a* or *nrxn3b* mutants (Fig S3D). Despite no change in unpaired postsynapses, both *nrxn3a* and *nrxn3b* single mutants exhibited a significant reduction in the total number of postsynapses (paired and unpaired) compared to controls (Fig S3D). Overall, this analysis indicates that loss of either Nrxn3a or Nrxn3b results in fewer complete synapses in mature lateral-line hair cells. In addition, this assessment suggests that zebrafish hair cells may rely on both Nrxn3a and Nrxn3b for proper pre-and post-synaptic pairing and synapse organization.

As each single mutant exhibited modest defects in ribbon-synapse organization, we tested whether Nrxn3a and Nrxn3b have overlapping contributions by examining the organization of synapses in lateral-line hair cells of *nrxn3a; nrxn3b* double mutants at 5 dpf (Fig 2A-F). Importantly, *nrxn3a; nrxn3b* neuromasts were grossly normal, and the number of hair cells per neuromast was unchanged (Fig 2G). However, we observed a ∼60% reduction in the number of complete ribbon synapses per hair cell in *nrxn3a; nrxn3b* mutants compared to controls (Fig 2A-F,H; synapses per hair cell, control: 3.19 ± 0.09, *nrxn3a;nrxn3b*: 1.3 ± 0.09, p < 0.0001). We also observed a dramatic increase in the number of unpaired pre- and post-synapses per cell in *nrxn3a; nrxn3b* mutants compared to control (Fig 2I-J). This result is especially striking for postsynapses, as we observed an increase in unpaired postsynapses despite a significant decrease in the total number (paired and unpaired) of postsynapses per cell (Fig 2J). In contrast, we found that the total number of presynapses per cell was unchanged (Fig 2I). Overall, this indicates that in mature hair cells, loss of Nrxn3 results in a dramatic decrease in ribbon-synapse numbers and a disruption in pre- and post-synaptic pairing.

**Fig 2.**
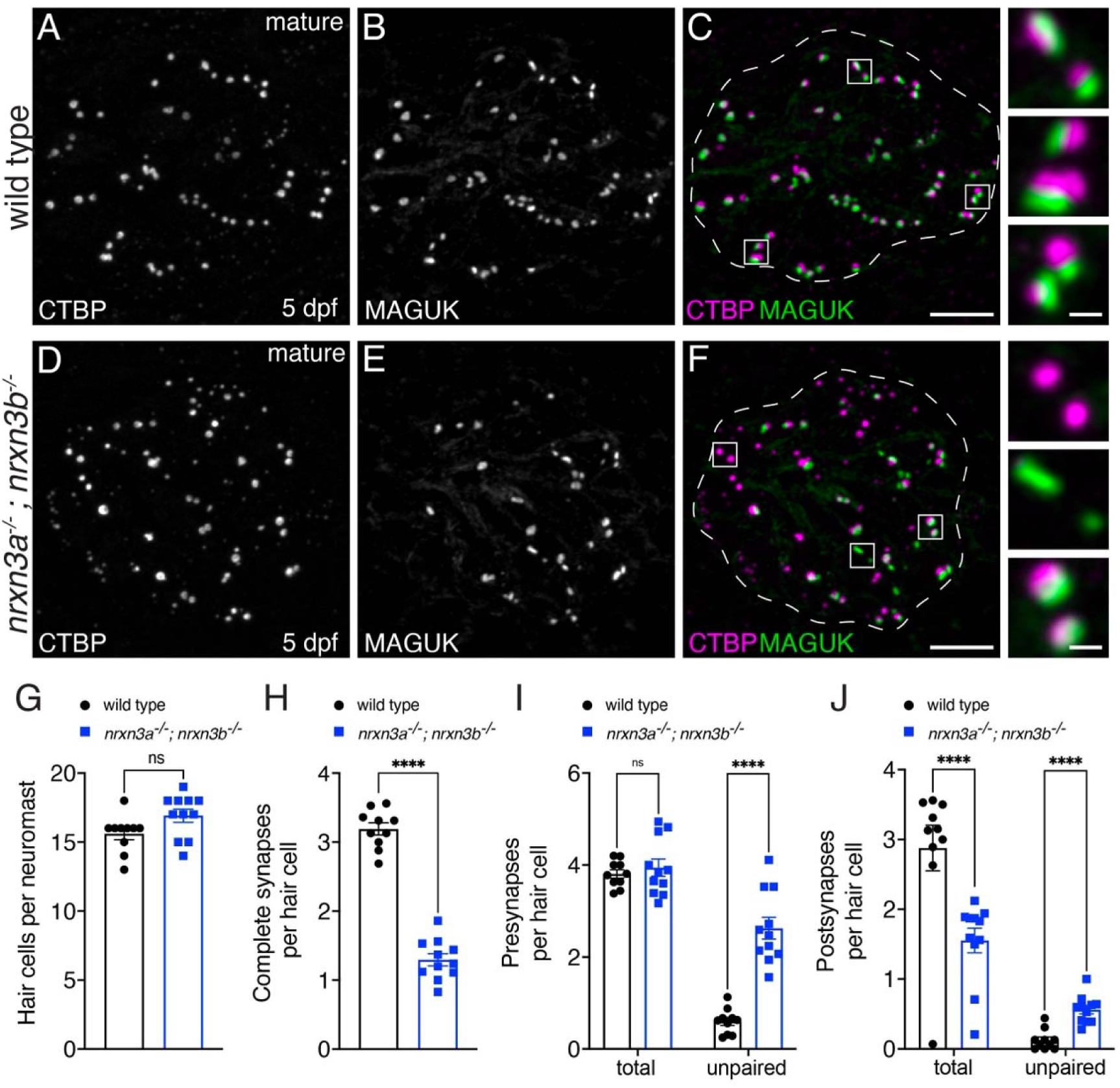
Loss of Nrxn3 reduces synapse numbers and pre- and postsynaptic pairing in mature lateral-line hair cells. (**A-F**) Confocal images of mature neuromasts (5 dpf) from wild-type controls (A-C) and *nrxn3a; nrxn3b* mutants (D-F). CTBP is used to label the presynapses (A,D), and MAGUK is used to label the postsynapses (B,E). Merged images are shown in C and F. The insets to the side in C and F show 3 examples of individual synapses. (**G-J**) Quantification reveals that wild-type controls and *nrxn3a; nrxn3b* mutants have a similar number of hair cells per neuromast (G). There are significantly fewer complete synapses per hair cell in *nrxn3a; nrxn3b* mutants compared to wild-type controls (H). Along with fewer complete synapses, there are significantly more unpaired presynapses (I) and postsynapses (J) per hair cell in *nrxn3a; nrxn3b* mutants compared to wild-type controls. While there is no change in the total number of presynapses (I), the total number of postsynapses (J) per hair cell is decreased in *nrxn3a; nrxn3b* mutants compared to wild-type controls. N = 10 wild-type and 11 *nrxn3a; nrxn3b* mutant neuromasts at 5 dpf. Synapse quantifications were replicated in at least three separate experiments. An unpaired t-test (2-tailed) was used in G and H, and a 2-way ANOVA was used in I and J. ns P > 0.05, ****P < 0.0001. Scale bars = 5 µm in C, 0.5 µm in the inset in C.

Within lateral-line neuromasts, there are hair cells with two different orientations; these distinct populations detect fluid flow in two opposing directions (for example, in the primary neuromasts of the pLL, flow from anterior to posterior (A to P) and flow from posterior to anterior (P to A); see Fig S4A). Because neurexins are also implicated in synapse selectivity (Gomez et al., 2021; Südhof, 2017), we assessed whether the synaptic defects in *nrxn3a; nrxn3b* mutants were present in hair cells of a specific orientation. We found that synapse loss was the same across the two populations of hair cells in *nrxn3a; nrxn3b* mutants (Fig S4B). These results show that Nrxn3 is critical for ribbon-synapse organization in hair cells of both orientations within the lateral line.

We also extended our synapse analysis of *nrxn3a; nrxn3b* mutants to hair cells in the inner ear. We examined hair cells in the zebrafish utricle (balance organ) and the medial crista (one of three organs detecting angular acceleration) (example images, Fig S5A-D). Within both inner-ear epithelia, we found that there were significantly fewer complete synapses per hair cell and significantly more unpaired presynapses and postsynapses in *nrxn3a; nrxn3b* mutants compared to controls (Fig S6). Overall, our analyses demonstrate that in zebrafish, both *nrxn3a* and *nrxn3b* are expressed in hair cells. In addition, Nrxn3a and Nrxn3b are essential for pre- and post-synaptic pairing and ribbon-synapse organization in mature hair cells of the lateral line and inner ear.

### Nrxn3 is required early in zebrafish hair-cell synapse assembly

Our results show that loss of Nrxn3 can dramatically impact the pairing of synaptic components in hair cells (Figs 2, S5, S6). For these initial analyses, we examined zebrafish hair cells at 5 dpf, a stage where the majority of hair cells are mature. Because hair cells are already mature at this stage, it is difficult to interpret whether Nrxn3 is required early to assemble synapses, or later in development to maintain or refine synapse numbers. Therefore, we examined hair cells in zebrafish at 3 dpf, a stage when hair cells are still developing, and sensory systems are not yet functional.

Using immunostaining, we quantified the number of complete synapses and unpaired pre- and post-synapses in developing hair cells at 3 dpf in *nrxn3a; nrxn3b* mutants (example images, Fig 3A-F). We found that developing hair cells in *nrxn3a; nrxn3b* mutants showed a significant, ∼35% reduction in the number of complete ribbon synapses per hair cell compared to controls (Fig 3G-H). Although significant, this reduction in complete synapses is less dramatic than the ∼60% reduction observed in mature hair cells (Fig 2). In developing hair cells, we also observed a dramatic increase in the number of unpaired presynapses (Fig 3I) and a more modest increase in unpaired postsynapses (Fig 3J). In contrast to mature hair cells at 5 dpf, in developing hair cells we did not observe a reduction in the total number of postsynapses per hair cell (Fig 3J). More complete synapses and a normal number of postsynapses at 3 dpf suggests that there may be synapses that are able to initially pair, but later fall apart, in *nrxn3a; nrxn3b* mutants. Overall, this analysis indicates that Nrxn3 plays an early role in synapse assembly but may also play a later role in synapse maintenance in lateral-line hair cells.

**Fig 3.**
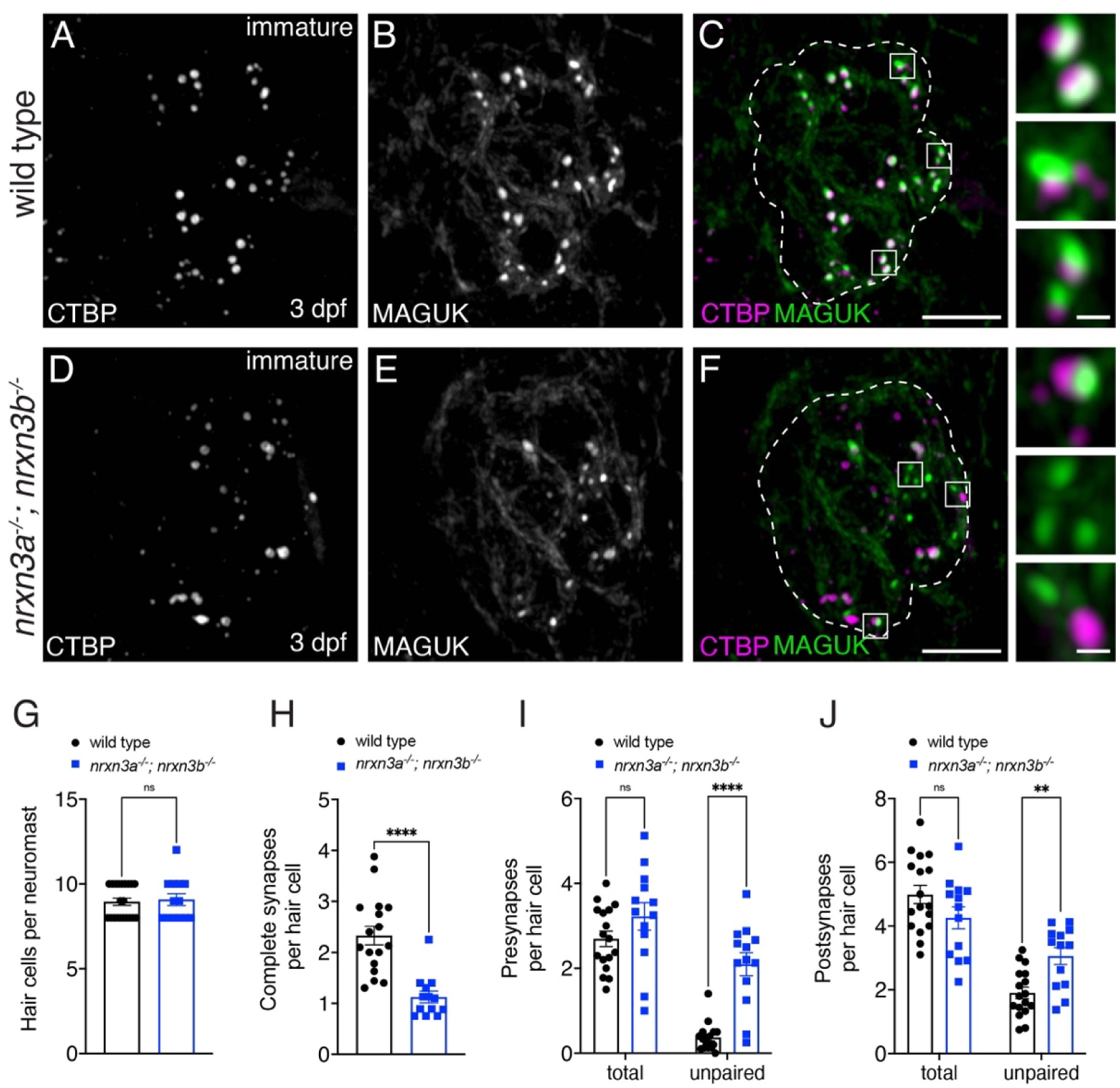
Nrxn3 is required for early for synapse assembly in lateral-line hair cells. (**A-F**) Confocal images of developing hair cells (3 dpf) from wild-type controls (A-C) and *nrxn3a; nrxn3b* mutants (D-F). CTBP is used to label the presynapses (A,D), and MAGUK is used to label the postsynapses (B,E). Merged images are shown in C and F. The insets to the side in C and F show 3 examples of individual synapses. Dashed lines in C and F outline the hair-cell region in each image. (**G-J**) Quantification reveals that wild-type controls and *nrxn3a; nrxn3b* mutants have a similar number of hair cells per neuromast (G). There are significantly fewer complete synapses per hair cell in *nrxn3a; nrxn3b* mutants compared to wild-type controls (H). Along with fewer complete synapses, there are significantly more unpaired presynapses (I) and postsynapses (J) per hair cell in *nrxn3a; nrxn3b* mutants compared to wild-type controls. In developing hair cells there is no change in the total number of presynapses (I), or postsynapses (J) per hair cell in in *nrxn3a; nrxn3b* mutants compared to wild-type controls. N = 17 wild-type and 13 *nrxn3a; nrxn3b* mutant neuromasts. Synapse quantifications were replicated in three separate experiments. An unpaired t-test (2-tailed) was used in G and H, and a 2-way ANOVA was used in I and J. ns P > 0.05, *P < 0.05, ****P < 0.0001. Scale bars = 5 µm in C and F, 0.5 µm in the insets.

### Nrxn3 alters pre- and post-synapse size and Ca**_V_**1.3 channel distribution in zebrafish

Studies in the central nervous system have found that loss of neurexins can alter the morphology and clustering of pre- and post-synaptic components. For example, previous work in neurons has shown that loss of neurexins can alter presynapse morphology and size, and impact the clustering of presynaptic calcium channels (Brockhaus et al., 2018; Luo et al., 2020; Uemura et al., 2022). Therefore, we next examined the morphology of synaptic components, including presynapses, postsynapses and Ca_V_1.3 channels, in *nrxn3a; nrxn3b* mutants at 5 dpf. We examined maximum intensity projections and quantified the 2D area and average intensity of these synaptic components. For our analysis, we examined the size of paired and unpaired synaptic components separately. We predicted that if Nrxn3 is important for synaptic organization, loss of Nrxn3 might specifically impact the architecture of paired synapses.

We first examined how loss of Nrxn3 impacts the average area (size) of presynaptic puncta (CTBP). We found that the average size of paired presynapses (paired CTBP-MAGUK puncta) was significantly increased in *nrxn3a; nrxn3b* mutants compared to controls (Fig 4A). We then examined the size of the unpaired presynapses in *nrxn3a; nrxn3b* mutants, and found they were a similar size compared to controls (Fig 4B). We also examined how loss of Nrxn3 impacts that size of postsynaptic puncta (MAGUK). We found that the average size of paired postsynapses (paired CTBP-MAGUK puncta) were significantly larger in *nrxn3a; nrxn3b* mutants, while unpaired postsynapses were a similar size compared to controls (Fig 4C,D). Altogether, this analysis indicates that in addition to a decrease in the number of complete synapses, *nrxn3a; nrxn3b* mutants form larger pre- and post-synapses compared to controls. This suggests that synapses do not properly coalesce in *nrxn3a; nrxn3b* double mutants, even when properly paired.

**Fig 4.**
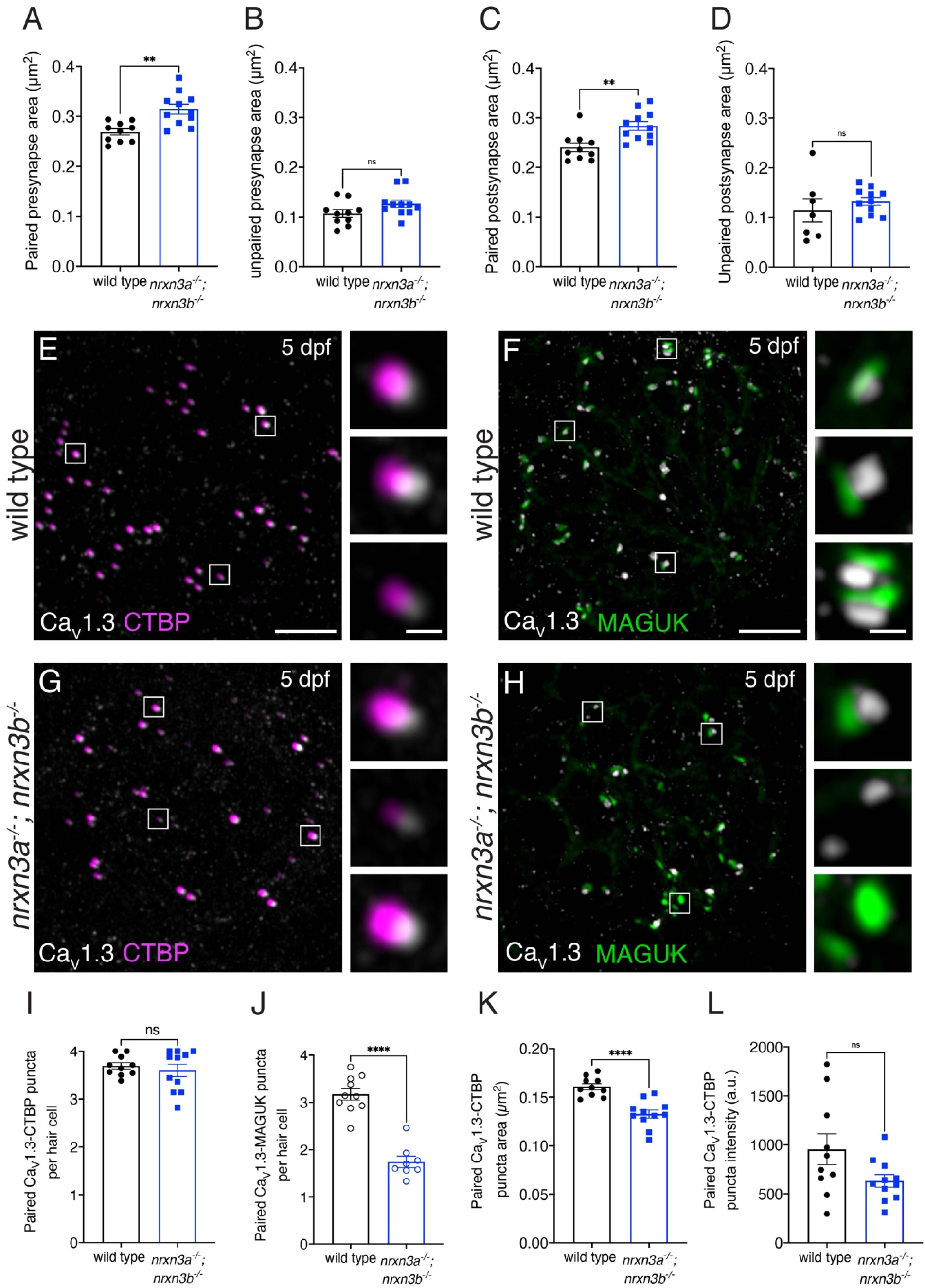
Loss of Nrxn3 impacts pre- and post-synapse size and Ca_V_1.3 channel localization in lateral-line hair cells. **(A-D)** There is a significant increase in the area of paired (A,C) but not unpaired (B,D) pre-and post-synapses in *nrxn3a; nrxn3b* mutants compared to wild-type controls. (**E-H**) Confocal images of mature neuromasts (5 dpf) from wild-type controls (E,F) and *nrxn3a; nrxn3b* mutants (G,H). An immunostain for CTBP is used to label the presynapses along with Ca_V_1.3 (E,G), or an immunostain for MAGUK is used to label the postsynapse and Ca_V_1.3 (F,H). Merged images are shown in E-H. The inset to the right of each merged image shows 3 examples of individual synapses. (**I-L**) Quantification reveals that the number of Ca_V_1.3-CTBP paired puncta per hair cell is the same in *nrxn3a; nrxn3b* mutants compared to wild-type controls (I). Compared to wild-type, *nrxn3a; nrxn3b* mutant neuromasts have a dramatically fewer Ca_V_1.3-MAGUK paired puncta per hair cell (J). The area (K) but not the average intensity (L) of Ca_V_1.3 puncta associated with CTBP puncta are significantly lower in *nrxn3a; nrxn3b* mutants compared to wild-type controls. N = 10 wild-type and 11 *nrxn3a; nrxn3b* mutant neuromasts in A-D and I, K and L and n = 10 wild-type and 8 *nrxn3a; nrxn3b* mutant neuromasts in J. Images and quantification are from larvae at 5 dpf. Ca_V_1.3 labeling was confirmed in two separate experiments. An unpaired t-test (2-tailed) was used in A-D and I-L. ns P > 0.05, **P < 0.01, ****P < 0.0001. Scale bars = 5 µm in E-H, 1 µm in the inset in E-H.

Previous work has shown that in hair cells, presynaptic Ca_V_1.3 channel distribution is shaped by presynapse size (Sheets et al., 2017). In our present study, we found that presynapse area and number was altered in *nrxn3a; nrxn3b* mutants. Therefore, we examined the number and distribution of Ca_V_1.3 channel puncta in *nrxn3a; nrxn3b* using an immunostain to label presynapses (CTBP) and Ca_V_1.3 channels. We first examined the number of presynapses paired with Ca_V_1.3 puncta (example images, Fig 4E,G). We found a similar number of CTBP-Ca_V_1.3 paired puncta per hair cell in *nrxn3a; nrxn3b* mutants compared to controls (Fig 4I). We then examined the number of postsynapses paired with Ca_V_1.3 puncta using an immunostain to label postsynapses (MAGUK) and Ca_V_1.3 channels (example images, Fig 4F,H). Here we found a reduced number of MAGUK-Ca_V_1.3 paired puncta per hair cell in *nrxn3a; nrxn3b* double mutants compared to controls (Fig 4J). This latter reduction mirrors the reduction in complete synapses in *nrxn3a; nrxn3b* mutants (Fig 2H). Our examination of Ca_V_1.3 pairing indicates that Nrxn3 is not required for presynapses to couple with Ca_V_1.3 channels. Instead Nrxn3 may be required to pair a presynapse and its associated Ca_V_1.3 channels to an adjacent postsynapse.

Lastly, we examined the size and distribution of Ca_V_1.3 channels within each CTBP-Ca_V_1.3 paired puncta. We found that the area of Ca_V_1.3 puncta was significantly reduced in *nrxn3a; nrxn3b* double mutants compared to controls (Fig 4K). Although we observed smaller Ca_V_1.3 puncta, the average intensity of each Ca_V_1.3 punctum was unchanged in *nrxn3a; nrxn3b* mutants compared to controls (Fig 4L). This indicates that on average, fewer Ca_V_1.3 channels may reside within each Ca_V_1.3 puncta in *nrxn3a; nrxn3b* mutants compared to controls. Together, this analysis of synapse morphology demonstrates that in addition to proper pre- and post-synapse pairing, Nrxn3 is important to establish proper pre- and post-synapse size.

### Nrxn3 plays a conserved role in synapse organization in mice

Many of the core genes required at hair-cell synapses are conserved between zebrafish and mammals (Sheets et al., 2021). Like zebrafish hair cells, scRNAseq studies in mice have shown that *Nrxn3* is expressed in hair cells in both the auditory and vestibular system (Cai et al., 2015; Elkon et al., 2015; Kolla et al., 2020). However, a role for NRNX3 in mammalian hair-cell synapse organization has not yet been demonstrated.

To determine whether Nrxn3 is required for synapse assembly in mice we used a conditional inactivation strategy. The *Nrxn3^flox^* strain (Aoto et al., 2015) was bred with the *Atoh1-Cre* driver (Matei et al., 2005) to abrogate both the α- and β-isoforms of NRXN3 in postmitotic hair cells. We examined ribbon synapses in auditory inner hair cells (IHCs) of *Atoh1-Cre; Nrxn^flox/flox^* mutants (*Nrxn3* mutants) and control animals by immunolabeling both presynapses (CTBP2) and postsynapses (GluR2). We examined IHCs at 4 (P28) and 6 (P42) weeks of age. At both ages we found a significant reduction (20-25%) in the number of complete synapses (paired CTBP2-GluR2 puncta) per IHC in 3 distinct tonotopic regions of the mouse cochlea (apex, mid and base thirds) in *Nrxn3* mutants compared to controls (Fig 5A-E, Fig S7A-E). These results indicate that NRXN3 plays a conserved role in synapse organization in mouse IHCs.

**Fig 5.**
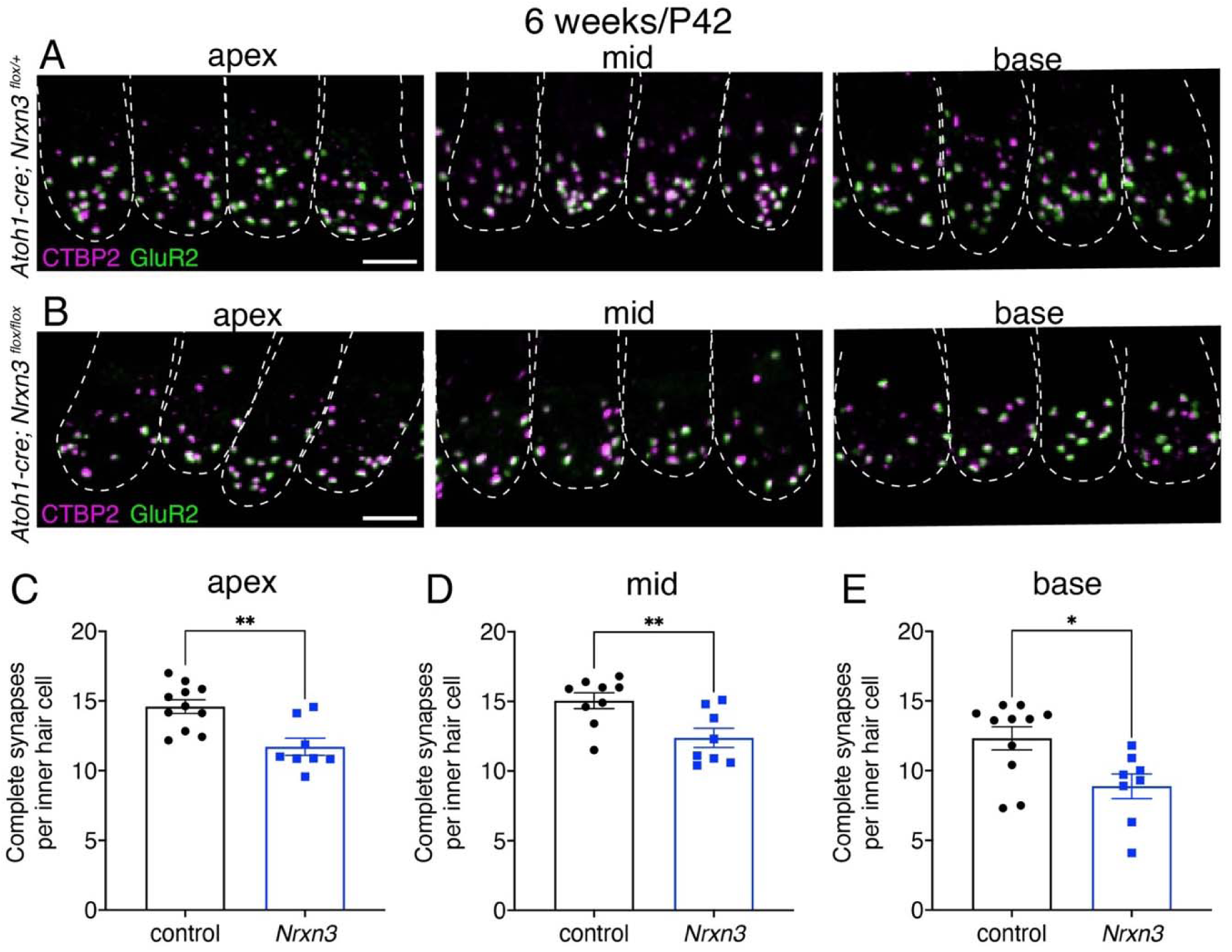
NRXN3 is required at 6 weeks for proper synapse number in mouse auditory inner hair cells. (**A-B**) Confocal images of mouse IHCs at 6 weeks (P42) from control (A) and *Nrxn3* mutant animals (*Atoh1-Cre; Nrxn3^flox/flox^*) (B). CTBP2 is used to label the presynapses (magenta), and GluR2 is used to label the postsynapses (green). Merged images show 4 IHCs from 3 different regions of the cochlea (apex, middle, basal thirds) for each genotype. Dashed lines indicate the outlines of hair-cell bodies in each image. (**C-E**) Quantification reveals that compared to controls, *Nrxn3* mutants have significantly fewer complete synapses per IHC at the apex (C) mid (D) and base (E). N = 70 control and 58 *Nrxn3* IHCs for the apex region, 70 control and 64 *Nrxn3* IHCs for the for mid region, 75 control and 57 *Nrxn3* IHCs for the for base region. These findings were compiled from 4 animals from each genotype and from 2 independent liters and immunostains. An unpaired t-test (2-tailed) was used in C-E. *P < 0.05, **P < 0.01. Scale bar = 5 µm in A.

### Nrxn3 disrupts ribbon-synapse function in hair cells in zebrafish

After verifying that Nrxn3 is essential for proper synapse organization in both zebrafish and mice, we next assessed the functional impact of fewer synapses. Previous work from our group established assays to assess hair-cell function in living zebrafish (Lukasz and Kindt, 2018). These assays rely on transgenic fish expressing GCaMP6s in hair cells or afferent neurons of the lateral-line system (Zhang et al., 2018). A fluid jet is used to stimulate hair cells, and GCaMP6s-dependent calcium signals are imaged during stimulation. To assay mechanotransduction and presynaptic calcium responses, we used a transgenic line expressing a membrane-localized GCaMP6s (memGCaMP6s) in lateral-line hair cells. We imaged calcium signals both at the mechanosensory hair bundles and at the presynapse (Fig 6A; Fig S8A (Zhang et al., 2018)). To assay postsynaptic activity, we used a transgenic line expressing GCaMP6s in afferent neurons and imaged calcium signals in the afferent terminals beneath lateral-line hair cells (Fig 6A (Zhang et al., 2018)).

**Fig 6.**
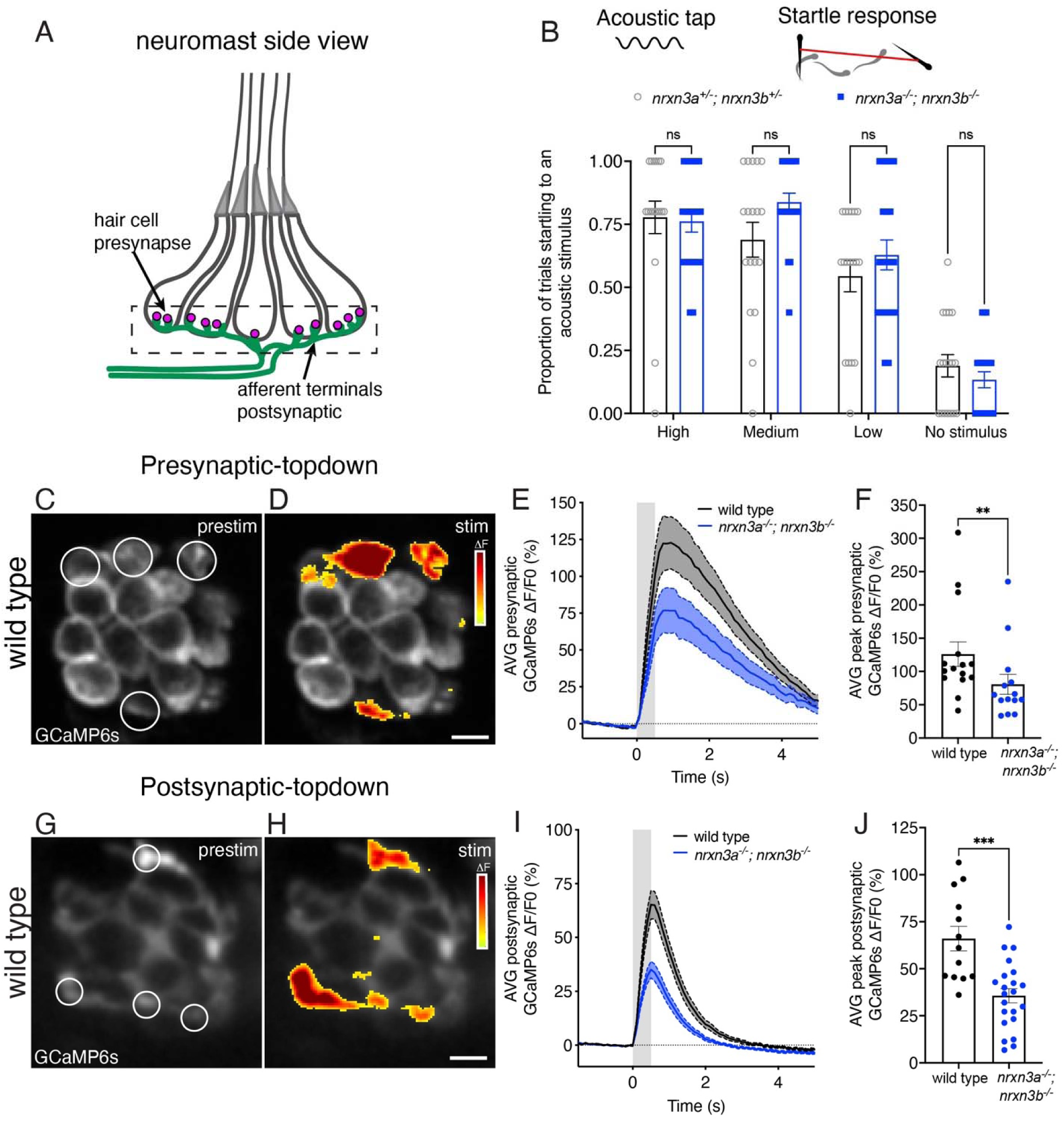
Nrxn3 is required for proper hair-cell synapse function in the lateral line. (**A**) Schematic of a neuromast shown from the side. The pre- and post-synaptic region used to measure GCaMP6s responses is indicated with a dashed box. (**B**) A vibrational acoustic tap stimulus was used at three stimuli of decreasing intensity to trigger an escape response in *nrxn3a^+/-^; nrxn3b^+/-^* double heterozygotes and *nrxn3a^-/-^; nrxn3b^-/-^* double mutants. The proportion of times (out of 5 trials) an animal responded to each stimulus is shown. N = 18 *nrxn3a; nrxn3b* double heterozygotes, and 21 *nrxn3a; nrxn3b* double mutants at 5 dpf. Behavioral data was acquired from two independents clutches and experimental days. (**C-D**) ΔF heatmaps show spatial patterns of presynaptic GCaMP6s increases in hair cells before (C) and during (D) a 500 ms fluid-jet stimulation in a wild-type neuromast. ROIs indicate synaptically active hair cells and examples of regions used to measure the average response per neuromast. (**E**) ΔF/F0 GCaMP6s traces showing average presynaptic GCaMP6 response during stimulation for wild-type controls (black) and *nrxn3a; nrxn3b* mutants (blue). Traces are displayed as mean, dashed lines are SEM, shaded gray represents the timing of the stimulus. (**F**) Maximum ΔF/F0 presynaptic calcium responses to stimulation for wild-type controls (black) and *nrxn3a; nrxn3b* mutants (blue). N = 15 wild-type and 14 *nrxn3a; nrxn3b* mutant neuromasts at 5-6 dpf. (**G-H**) ΔF heatmaps show spatial patterns of postsynaptic GCaMP6s increases in the afferent terminal before (G) and during (H) a 500 ms fluid-jet stimulation in a wild-type neuromast. ROIs indicate synaptically active terminals and examples of regions used to measure the average active postsynaptic response per neuromast. (**I**) ΔF/F0 GCaMP6s traces showing average postsynaptic GCaMP6 response during stimulation for wild-type controls (black) and *nrxn3a; nrxn3b* mutants (blue). Traces are displayed as mean, dashed lines are SEM. (**J**) Maximum ΔF/F0 postsynaptic calcium responses to stimulation for wild-type controls (black) and *nrxn3a; nrxn3b* mutants (blue). N = 13 wild-type and 22 *nrxn3a; nrxn3b* mutant neuromasts at 4-5 dpf. Each dot in F and J represents the average response from a single neuromast. Calcium imaging findings were acquired from two independents clutches and experimental days. A two-way ANOVA was used in B, a Mann-Whitney test in F and an unpaired t-test was used in J. ns P > 0.05, ***P < 0.001, **P < 0.01. Scale bars = 5 µm in D and H.

Using GCaMP6s in hair cells, we first assessed whether *nrxn3a; nrxn3b* mutants have normal mechanotransduction (i.e. normal ability to detect sensory stimuli). We found that the magnitude of GCaMP6s signals measured in hair bundles was not significantly different in *nrxn3a; nrxn3b* mutants compared to controls (Fig S8B-C). After verifying normal mechanotransduction, we next assessed the magnitude of evoked GCaMP6s signals measured in the presynaptic region of hair cells. We found that the magnitude of presynaptic GCaMP6s signals was reduced by ∼35% in *nrxn3a; nrxn3b* mutants compared to controls (Fig 6C-F and Fig S9A-F). These signals were measured at the level of individual hair cells, not individual synapses. Thus, a reduction in presynaptic calcium signals could be due to fewer complete synapses, smaller presynapses, altered calcium channel density, or some combination of these possibilities.

It is important to understand how changes in presynaptic calcium activity are reflected in the afferent neurons, as afferents encode information that is ultimately carried to the downstream circuitry. Therefore, we next examined evoked GCaMP6s signals in the afferent terminals of *nrxn3a; nrxn3b* mutants. We observed a ∼50% reduction in the magnitude of evoked GCaMP6s signals in the terminals of *nrxn3a; nrxn3b* mutants compared to controls (Fig 6G-J and Fig S9G-L). Overall, our calcium imaging experiments indicate that the synapse loss in *nrxn3a; nrxn3b* mutants result in a dramatic reduction in afferent terminal responses.

After assessing how synapse loss in *nrxn3a; nrxn3b* mutants impacts synapse function, we next examined how these deficits impacted behavior. One robust behavioral assay to assess hair-cell function in zebrafish is the acoustic startle response. In response to an acoustic/vibrational stimulus, zebrafish will exhibit a stereotyped startle response (Kimmel et al., 1974). This behavior relies on hair cells in both the inner ear and the lateral line and is appropriate to use because we observed synapse loss in *nrxn3a; nrxn3b* mutants in both sensory systems (Figs 2, Fig S5, S6). For an in-clutch comparison, we compared *nrxn3a; nrxn3b* double mutants to *nrxn3a^+/-^; nrxn3b^+/-^* double heterozygotes. We assayed the acoustic startle response using an automated Zantiks behavioral system at three different intensities. Using this approach, we observed no difference in the proportion of animals startling at any stimulus intensity between the two genotypes (Fig 6B). A surprisingly normal acoustic startle response in *nrxn3a; nrxn3b* mutants shows that a dramatic reduction in synapse numbers is not sufficient to impact this behavior. It remains possible that other lateral-line-mediated behaviors are more sensitive to reductions in hair-cell synapse number. However, our results broadly suggest that hair-cell sensory systems are extremely robust and may not require all synapses to function.

## Discussion

In zebrafish and mice, we find a dramatic reduction in the number of ribbon synapses in hair cells when Nrxn3 is absent (mouse Nrxn3 cKO and *nrxn3a; nrxn3b* zebrafish mutants). In zebrafish, we show that presynapses are present in normal numbers and are still tightly coupled to Ca_V_1.3 channels in *nrxn3* mutants, but they fail to pair with postsynapses. Further, we demonstrate that both pre- and post-synaptic calcium responses are reduced in the absence of Nrxn3. Overall, this work highlights a conserved role for Nrxn3 in the organization of ribbon synapses in hair cells.

### Postsynaptic partners for Nrxn3 at the ribbon synapse in hair cells

Our work demonstrates that Nrxn3 is required in zebrafish and mouse hair cells for proper organization of ribbon synapses. Neurexins serve as presynaptic receptors for several extracellular binding partners to facilitate synapse assembly. For α-neurexins these partners include: secreted cerebellins and neuroexophilins, as well as transmembrane proteins such as neuroligins, Dystroglycan, leucine-rich repeat transmembrane proteins (LRRTM), and Calsyntenin-3 (Boucard et al., 2005; Dai et al., 2022; Hauser et al., 2022; Kim et al., 2020; Ko et al., 2009; Sugita et al., 2001; Trotter et al., 2023). Recent work in mice has demonstrated a role for α-neurexin binding partners NLGN1 and NLGN3 at the postsynapses of mouse auditory IHCs (Ramirez et al., 2022). When NLGN1 and NLGN3 are lost, there is a ∼25% loss of ribbon synapses in IHCs (Ramirez et al., 2022). In our work, we also observed a ∼20-25% reduction in ribbon synapses in *Nrxn3* mouse mutants (Fig 5, Fig S7). Based on these results, it is possible that α-Nrxn3 is a presynaptic binding partner for NLGN1 and NLGN3 at auditory IHCs synapses in mice. In the future, it will be interesting to examine zebrafish mutants lacking NLGN1 and NLGN3 to understand whether this complex plays a conserved role in vertebrate species. In addition, it will be interesting to examine both mice and zebrafish lacking both Nrxn3 and Nlgn1/3 to understand the requirements of this adhesion complex at ribbon synapses in more detail.

In mice lacking NLGN1 and NLGN3 or NRXN3 and zebrafish lacking α-Nrxn3 there is a clear reduction in the number of complete ribbon synapses–yet intact synapses remain. In our zebrafish *nrxn3* mutants, β-Nrxn3 is still present and could potentially compensate for the loss of α-Nrnx3. In contrast, our mouse *Nrxn3* mutant lacks both α- and β-Nrxn3 (Aoto et al., 2015). Therefore, in mice neither NRXN3 nor NLGN1/3 are strictly required to organize all ribbon synapses in IHCs. Whether β-Nrxn3 (in zebrafish) or other neurexins (in zebrafish or mice) can partially compensate when NRXN3 is lost, or whether these remaining synapses simply rely on a completely different synaptic adhesion complex, remains to be determined.

### A role for Nrxn3 in hair-cell synapse specificity?

If Nrxn3 and Nlgn1/3 are not required to organize all ribbon synapses in mice or zebrafish, it is possible that this complex is only required for a specific subset of synapses. One example of subset specificity in the lateral line is selective innervation based on the orientation of hair cells (Nagiel et al., 2008). Here, afferent neurons innervate lateral-line hair cells based on the direction of fluid that they sense (for example, anterior or posterior flow in the primary posterior-lateral line, Fig S4, (López-Schier and Hudspeth, 2006)). In our work, we observed a similar loss of ribbon synapses in hair cells that sense anterior and posterior fluid flow (Fig S4B). This indicates that α-Nrnx3 is not required for selective innervation based on hair-cell orientation.

Another level of innervation specificity in the lateral line is reflected in the wiring pattern each neuron makes within the afferent terminals beneath neuromast hair cells. Numerous studies have demonstrated that in the lateral line, hair cells are redundantly innervated by multiple neurons (Haehnel et al., 2012; Nagiel et al., 2008). More recent connectomic work has shown that each neuromast contains a dominant afferent neuron that innervates nearly all hair cells (one for each orientation); these neurons form 75% of the synapses (Dow et al., 2018). In addition to these dominant neurons, 1-3 additional afferent neurons form the remaining synapses within the neuromast. Therefore, it is possible that in the lateral line, □-Nrnx3 is required specifically to form synapses made by the dominant neurons. This is consistent with the 60% loss in synapses we observed in zebrafish *nrxn3* mutants. In this scenario, a separate synaptic adhesion complex could be used to form synapses in the less dominant neurons. This would suggest that there may be distinct subsets of neurons in the posterior-lateral line. While studies have shown that there are both functionally (spike rate) and morphologically (innervation pattern) distinct neurons within the lateral-line nerve, currently there are no molecular markers to distinguish these subtypes (Dow et al., 2018; Liao and Haehnel, 2012). In the future, it will be important to investigate the role Nrxn3 plays in the lateral line with regards to these potential subtypes.

While neuronal subtypes are less defined in the lateral line, in the mouse auditory system, molecular studies (scRNAseq) have demonstrated that there are at least three main subtypes of afferent neurons (Type I spiral ganglion neurons) that innervate auditory IHCs within the mouse cochlea (Shrestha et al., 2018; Sun et al., 2018). Each subtype synapses onto hair cells at roughly distinct spatial locations (across the pillar and modiolar faces of the cell) and likely corresponds to a distinct functional profile (Liberman et al., 2011; Petitpré et al., 2018; Taberner and Liberman, 2005). Work on NLGN1 and NLGN3 in the mouse cochlea has shown that NLGN1 is localized to postsynapses contacting the modiolar face of the auditory IHCs (Ramirez et al., 2022). In contrast, the pillar face of auditory IHCs is populated by postsynapses containing NLGN3 or NLGN1/3. Whether a particular combination of NRXN3 and NLGN1 or 3 are required to form synapses at distinct spatial locations on mouse auditory IHCs remains to be determined.

### Nrxn3 Interactions with core presynaptic components

Although neurexins are considered classic synaptic adhesion molecules, genetic loss of neurexins does not confer synapse loss in all contexts. For example, in the brainstem of *α-Nrxn1/2/3* triple KO mice, there is no loss of glutamatergic synapses (Missler et al., 2003). Instead, Missler et al. found that α-NRXNs play a key role in calcium-mediated neurotransmission; this work found that presynaptic calcium channel function was reduced despite a normal number of synapses and channels. These data led to the hypothesis that α-Nrxns may act to couple calcium channels to presynaptic machinery. In a separate study, *α-Nrxn1/2/3* triple KO mice were shown to impact presynaptic Ca_V_2.1 channel clustering at a central auditory synapse, the calyx of Held (Luo et al., 2020). This study also found that loss of α-NRXNs did not result in synapse loss. Instead, mutant synapses contained fewer Ca_V_2.1 channels. These defects ultimately led to impaired neurotransmission.

In zebrafish hair cells, we found that loss of Nrxn3 also impacts Ca_V_1.3 channels at the hair-cell presynapse. Our work on zebrafish *nrxn3* mutants found smaller clusters of the α-subunit of Ca_V_1.3 channels (Fig 4E-H, K), and smaller presynaptic calcium responses in the hair cells of the lateral line (Fig 6E-F). In hippocampal neurons, α-NRXNs can regulate presynaptic calcium responses via interactions with α2δ1 auxiliary subunits of Ca_V_2.1 channels (Brockhaus et al., 2018). Based on this work, it is possible that the reduced presynaptic calcium signal that we observed in *nrxn3* mutants could result, at least in part, from loss of interactions between Nrxn3 and an α2δ subunit of Ca_V_1.3 channels in hair cells. Other studies have found that α2δ subunits act as trans-synaptic organizers of glutamatergic synapses by aligning the pre- and post-synaptic active zones (Schöpf et al., 2021). Consistent with this idea, work in mouse auditory IHCs has shown that α2δ2 subunits are required for proper gating properties of Ca_V_1.3 channels as well as alignment of these channels with the postsynapse (Fell et al., 2016). In future work, it will be interesting to pursue both the physical and functional link between Nrxn3 and all subunits of Ca_V_1.3 channels.

The most straightforward way to understand the interaction between Nrxn3 and other synaptic components is to visualize Nrxn3 localization. Unfortunately, neurexin molecules are notoriously difficult to label either using immunohistochemistry or via tagged proteins. In the future it will be important to generate endogenously tagged Nrxn3 animal models to visualize whether Nrxn3 is present at ribbons synapses in hair cells. In recent years, adding endogenous tags to proteins in mice and zebrafish has become more straightforward, making this approach more straightforward (Carrington et al., 2022; Morrow et al., 2021).

### Functional consequences of synapse loss in Nrxn3 mutants

In our study, we observed a dramatic loss of synapses in zebrafish hair cells in *nrxn3* mutants (Fig 2: 60% lateral line; Fig S6: 27% medial crista, 44% anterior macula). Along with this synapse loss, we also observed a dramatic decrease in the evoked pre- and post-synaptic calcium signals in hair cells of *nrxn3* mutants (Fig 6, 35% presynaptic and 45% postsynaptic reduction). Despite this decrease in synaptic function, *nrxn3* homozygotes were viable as adults and did not have any overt vestibular (circling behavior or difficultly remaining upright) or auditory defects (Fig 6B). This suggests that in zebrafish, there is enough redundancy built into hair-cell sensory systems that fewer synapses and decreased synaptic function do not affect these behaviors. This result it in line with work on auditory synaptopathy in mammals where a loss of IHC ribbon synapses does impact hearing thresholds, but instead is linked to more subtle hearing deficits (Liberman et al., 2016). Based on this work on auditory synaptopathy, it is possible that more subtle behavioral deficits exist in *nrxn3* zebrafish mutants. For example, *nrxn3* mutants may not be able to rheotax (use their lateral line to orient in a constant flow), or they may not be able to detect a particular auditory frequency or threshold. In the future, it will be interesting to study more complex hair-cell mediated behaviors in *nrxn3* zebrafish mutants in more detail.

In humans, *NRXN3* sequence variants and mutations have been linked to alcohol and drug abuse, obesity and autism (Heard-Costa et al., 2009; Hishimoto et al., 2007; Lachman et al., 2007; Vaags et al., 2012). Currently no studies have linked hearing loss in humans to the *NRXN3* locus, or any other *NRXN* locus. In mice, a genome-wide association study identified a locus near *Nlgn1* that was linked to compromised auditory brain-stem responses (Ramirez et al., 2022). In addition, *Nlgn1/3* mutant mice have impaired hearing and are more vulnerable to noise trauma. In the future it will be important to further examine *Nrxn3* mutant mice to investigate whether they have hearing and balance defects and whether they are more vulnerable to noise trauma.

Our work demonstrates that Nrxn3 is critical for synapse assembly in both mouse and zebrafish hair cells. However, the majority of work on the assembly of ribbon synapses in our current study and in other studies relies on information obtained from static images. It is important to note that development is not static but is instead dynamic and is best studied in living tissue over time. In future work, we will continue to use the zebrafish system–which is ideal for live imaging–to determine the exact role that Nrxn3 plays in synapse assembly *in vivo*. Overall, elucidating how hair-cell synapses assemble is essential to understand how to re-form synapses when they are lost after noise- or age-related hearing loss.

## Acknowledgements

The Zebrafish International Resource Center (ZIRC) provided the *nrxn3a^sa11330^* and *nrxn3b^sa36960^* ENU alleles used in this study. We thank the Reproductive Science group at The Jackson Laboratory for cryorecovery of the *Nrxn3^flox^* strain. This work was made possible in part by software funded by the NIH: Fluorender (VVDViewer) “An Imaging Tool for Visualization and Analysis of Confocal Data as Applied to Zebrafish Research”, R01-GM098151-01. We thank Drs. Katie Drerup and Candy Wong for their thoughtful comments on our manuscript.

## Data availability

All raw data for this work is posted on Dryad (link to be added upon completion).

## Contributions

AJ, ZCL, KK and NM performed immunohistochemistry and confocal imaging to examine *nrxn3* zebrafish mutants. BT did the mouse immunohistochemistry. ZCL wrote the code to quantify all immunohistochemistry and RNA FISH images. SD did the behavioral analyses. KP performed RNA FISH to examine *nrxn3* expression in zebrafish hair cells and assisted with genotyping and husbandry. EC and KK performed the calcium imaging for pre- and post-synaptic responses respectively. AJ, ZCL, EC and KK made figures and wrote the manuscript. All authors edited the manuscript.

## Diversity and Inclusive statement

We strive to make our group the most inclusive, diverse, and equitable space possible. We acknowledge the inequities that exist within STEM. We hold ourselves accountable and continuously recognize and work against these inequities. All scientists on this project had different levels of training, knowledge, personal circumstances, and backgrounds. Our lab seeks to provide scientists with an academic space that acknowledges, supports, and celebrates these differences.

## Materials and Methods

### Zebrafish strains and husbandry

Zebrafish (*Danio rerio*) were grown at 30°C using a 14 hr light, 10 hr dark cycle. Larvae were raised in E3 embryo medium (51mM NaCl, 0.171mM KCl, 0.331mM CaCl_2_, and 0.331mM MgSO_4_, pH 7.2). Zebrafish work performed at the National Institute of Health (NIH) was approved by the Animal Use Committee at the NIH under animal study protocol #1362-13. Larvae were examined at either 3 days post fertilization (dpf) or 5 dpf unless stated otherwise. The following previously established lines were used in this study: *myo6b:memGCaMP6s^idcTg1^* and *en.sill,hsp70l:GCaMP6s^idcTg8^* (Jiang et al., 2017; Zhang et al., 2018). In addition to these lines, two sanger mutants were obtained from the Zebrafish International Resource Center (ZIRC) and used in this study: *nrxn3a^sa11330^* and *nrxn3b^sa36960^*. The *nrxn3a^sa11330^* mutant results in a premature stop codon in the second LNS domain (C to stop at amino acid 455/1697 in the α isoform, ENSDART00000088179.5). This allele was genotyped using standard PCR and sequencing with the following primer sets: FWD: 5’-AATGAACTCTTTAAAAGGAGCA-3’ and REV: 5’-TCCACTTTTGTGTTCTTCTGGC-3’. The *nrxn3b^sa36960^* mutants results in a point mutation leading to a premature stop codon in the first LNS domain (R to stop at amino acid 135/1687 in the α isoform, ENSDART00000127050.3). This allele was genotyped using standard PCR and sequencing with the following primer sets: FWD: 5’-TCACTGGCACTTTGCTACAATC-3’ and REV: 5’-GTTGGAACCTTATTGCCGTAAC-3’. Each mutant line was outcrossed 3 times before use. After outcrossing, the *nrxn3a^sa11330^* and *nrxn3b^sa36960^* mutants were crossed to produce double mutants: *nrxn3a^-/-^; nrxn3b^-/^*^-^. For comparisons, *nrxn3a^-/-^,* and *nrxn3b ^-/-^,* and *nrxn3a^-/-^; nrxn3b^-/^*^-^ mutants were either compared to their respective wild-type siblings or to wild-type larvae collected and grown at the same time as double mutants. For behavioral experiments, *nrxn3a^+/-^; nrxn3b^+/-^* double heterozygotes were compared to double mutants obtained from the same clutch of embryos to directly compare siblings.

### Mouse strains and husbandry

The *Nrxn3^flox^* strain was cryorecovered at The Jackson Laboratory from stock JR#014157 (B6;129-*Nrxn3^tm3Sud/J^*; MGI:5437468) (Aoto et al 2015). In this strain, the first common exon for the α and β transcripts (exon 18) is flanked by *loxP* sites. The *Atoh1-Cre* driver used to inactivate *Nrxn3* in post-mitotic hair cells is stock JR#011104 (*B6.Cg-Tg(Atoh1-cre)1Bfri/J*; MGI:3775845) (Matei et al., 2005). *Atoh1-Cre; Nrxn3^flox/flox^* mutants were compared to control littermates of the following genotypes: *Atoh1-Cre; Nrxn3^flox/+^*(this control genotype is depicted in the Figures)*, Nrxn3^flox/flox^, Nrxn3^flox/+^*. Both males and females were included in the study. Animals were maintained under standard housing conditions (14h light/10h dark cycle, ambient temperature and normal humidity). All mouse work was reviewed for compliance and approved by the Animal Care and Use Committee of The Jackson Laboratory.

### Zebrafish immunohistochemistry and imaging

Immunohistochemistry was performed on whole larvae at either 3 dpf or 5 dpf. Whole larvae were fixed with paraformaldehyde (PFA 4%; Thermoscientific; 28906) in PBS at 4°C for 3.5 hr. For Ca_V_1.3 labeling (Ca_V_1.3, Otoferlin, MAGUK or Ca_V_1.3, Paravalbumin, CTBP), all wash, block and antibody solutions were prepared with PBS + 0.1% Tween (PBST). For pre- and post-synaptic labeling (rabbit anti-MYO7A, CTBP, MAGUK), all wash, block and antibody solutions were prepared with PBS1+11% DMSO, 0.5% Triton-X100, 0.1% Tween-20 (PBDTT). After fixation, larvae were washed 4 × 5 min in PBST or PBDDT. For Ca_V_1.3 labeling, prior to block, larvae were permeabilized with acetone. For this permeabilization, larvae were washed for 5 min with H_2_O in glass vials. The H_2_O was removed and replaced with ice-cold acetone and larvae placed at −20°C for 5 min, followed by a 5 min H_2_O wash. The larvae were then washed for 4 × 5 min in PBST. For all immunolabels, larvae were blocked overnight at 4°C in blocking solution (2% goat serum, 1% bovine serum albumin, 2% fish skin gelatin in PBST or PBDTT). After block, larvae were incubated in primary antibodies in antibody solution (1% bovine serum albumin in PBST or PBDTT) overnight, nutating at 4°C. The next day, the larvae were washed for 4 × 5 min in PBST or PBDTT to remove the primary antibodies. Secondary antibodies in antibody solution were added and larvae were incubated for 2 hrs at room temperature, with minimal exposure to light. Secondary antibodies were removed by washes with PBST or PBDTT for 4 × 5 min. Larvae were mounted on glass slides with Prolong Gold (ThermoFisher Scientific) using No. 1.5 coverslips.

Fixed samples were imaged on an upright LSM 980 laser-scanning confocal microscope with an Airyscan 2 attachment using Zen Blue 3.4 (Carl Zeiss) and a 63x/1.4 NA Plan Apo oil immersion objective lens. Z-stacks were acquired every 0.151µm with a 0.043 µm X-Y pixel size for lateral-line and medial-crista hair cells, and every 0.151µm with a 0.067 µm X-Y pixel size for hair cells in the anterior macula. The Airyscan z-stacks were autoprocessed in 2D. Experiments were imaged with the same acquisition settings to maintain consistency between comparisons. For presentation in figures, images were further processed using Fiji.

### Mouse immunohistochemistry and imaging

Temporal bones were isolated, and an insulin syringe was used to gently flush cold paraformaldehyde (PFA 4%; Electron Microscopy Sciences; 15710) through the cleared oval and round windows after poking a small hole at the cochlear apex. Temporal bones were then immersion-fixed in PFA for 1 hour at 4°C, washed in PBS, and rotated overnight in EDTA 4% for decalcification. The next day, cochleae were dissected in 3 approximate thirds (base, mid and apex) before blocking and permeabilization for 1 hr at room temperature under agitation (1% bovine serum albumin; 0.5% Triton X-100). The following primary antibodies were used: CTBP2, GluR2 and mouse anti-MYO7A. Primary and secondary antibodies were incubated overnight at 4°C in PBS. Samples were washed 3 times in PBS + 0.05% Triton X-100 after each antibody incubation and finally post-fixed in PFA 4% for at least 1 hr at room temperature. Samples were then mounted flat in Mowiol mounting medium (Calbiochem/MilliporeSigma 4759041) using two layers of office tape as a spacers for the coverglass (18x18mm #1.5). Mowiol (10% w/v) was prepared in (25% w/v) glycerol and 0.1M Tris-Cl pH8.5.

Mounted samples were imaged on an upright LSM 980 laser-scanning confocal microscope with using Zen Blue 3.4 (Carl Zeiss) and an 63x 1.4 NA oil objective lens. Z-stacks were acquired every 0.250□µm with an 0.085 µm X-Y pixel size in confocal mode. For presentation in figures, imaged were further processed using Fiji.

Primary antibody list.

*Primary antibody list*
| Antigen | Species; subtype | Concentration | Catalog or reference |
| --- | --- | --- | --- |
| Pan-MAGUK | Mouse; IgG1 | 1:500 | Millipore MABN7 |
| MYO7A | Rabbit | 1:1,000 | Proteus 25-6790 |
| MYO7A | Mouse; IgG1 | 1:400 (mouse)<br>1:1000 (zebrafish) | Developmental Study<br>Hybridoma Bank 138-1 |
| Otoferlin | Mouse; IgG2a | 1:1,000 | DSHB HCS-1 |
| Ca<sub>v</sub>1.3 | Rabbit | 1:1,000 | (Sheets et al., 2011) |
| Pan-CTBP | Mouse; IgG2a | 1:1,000 | Santa Cruz sc-3878 |
| GluR2 | Mouse; IgG2a | 1:100 | Millipore/Sigma MAB397 |
| CTBP2 | Rabbit | 1:400 | Synaptic Systems 192103 |

Secondary antibody and co-label list.

*Secondary antibody and co-label list*
| Antigen | Species; fluorophore | Concentration | Catalog or reference |
| --- | --- | --- | --- |
| Mouse IgG2a | Goat; Alexa 488, Alexa 555, Alexa 647 | 1:1,000 | ThermoFisher #A-21131, #A-2113, # A-21241 |
| Mouse IgG1 | Goat; Alexa 647, Alexa 555 | 1:1,000 | ThermoFisher #A-21240, #A-21422 |
| Mouse IgG | Donkey; Alexa 488 | 1:1,000 | ThermoFisher # A-21202 |
| Rabbit IgG | Donkey; Alexa 555 | 1:1000 | ThermoFisher #A-31572 |
| Rabbit IgG | Goat; Alexa 488, Alexa 555 | 1:1,000 | ThermoFisher #A-11008, # A-21428 |
| F-actin | Phalloidin Alexa 488 | 1:1000 | ThermoFisher #A-12379 |

### RNA FISH to detect *nrxn3a* and *nrxn3b* mRNA in lateral-line hair cells

To detect mRNA for *nrxn3a and nrxn3b* in zebrafish, we followed the Molecular Instrument-RNA FISH Zebrafish protocol, Revision Number 10 (https://files.molecularinstruments.com/MI-Protocol-RNAFISH-Zebrafish-Rev10.pdf), with a few minor changes to the preparation of fixed whole-mount larvae. For our dehydration steps, we dehydrated using the following methanol series: 25, 50, 75, 100, 100% methanol, with 5 min for each step in the series. To permeabilize, we treated larvae with 10 µg/mL proteinase K for 20 min. RNA FISH probes were designed to target the long α form of zebrafish *nrxn3a* and *nrxn3b* (Molecular Instrument Probe lot # PRP848, PRP849). After completing the RNA FISH protocol, we mounted the larvae in ProLong Gold Antifade (ThermoFischer, P36930) under 1.5 coverglass. Samples were imaged on an upright LSM 980 laser-scanning confocal microscope with an Airyscan 2 attachment using Zen Blue 3.4 (Carl Zeiss) and a 63x/1.4 NA Plan Apo oil immersion objective lens. Airyscan z-stacks were acquired every 0.1601µm with a 0.043 µm X-Y pixel size. The Airyscan z-stacks were autoprocessed in 2D. For presentation in figures, imaged were further processed using Fiji. RNA FISH label of *nrxn3a* and *nrxn3b* mRNA were further quantified in Fiji. The RNA FISH label of *nrxn3a* and *nrxn3b* mRNA in *nrxn3a; nrxn3b* double mutants was still present, but further quantification (see quantification information below) revealed a significant reduction in *nrxn3a* and *nrxn3b* mRNA (Fig S2A-G). A reduction in *nrxn3a* and *nrxn3b* mRNA suggests that these transcripts could undergo some nonsense mediated decay in *nrxn3a; nrxn3b* double mutants.

### Image processing and quantification of hair-cell synapses and RNA FISH

Z-stack image acquisitions from zebrafish and mouse confocal images were processed in Fiji (Schindelin et al., 2012). Researchers were blinded to genotype during analyses. In zebrafish neuromasts, hair-bundle orientation was scored manually (neuromasts L1-L4) relative to the midline of the muscle somites. Hair-cell numbers were counted manually based on MYO7a, Paravalbumin, or Otoferlin labeling. Prior to automated puncta quantification, each channel was background subtracted using rolling-ball radius background subtraction. Then each z-stack was max-intensity projected. A mask was generated by manually outlining the region or interest (ie: hair cells) in the reference channel. This mask was then applied to the z-projection of each synaptic component or RNA FISH channel.

An automated synapse quantification was then performed using a customized Fiji-based macro, “Complete Synapse Counter v5.2”. In this macro, each masked image was thresholded using an adaptive thresholding plugin by <u>Qingzong TSENG</u> (https://sites.google.com/site/qingzongtseng/adaptivethreshold) to generate a binary image of the puncta (presynaptic, postsynaptic, Ca_V_1.3 cluster or RNA FISH puncta). Individual synaptic or RNA FISH puncta were then segmented using the particles analysis function in Fiji. For particle analysis, the following minimum size thresholds were applied: zebrafish lateral-line images – CTBP: 0.025 μm^2^, MAGUK: 0.04 μm^2^, Ca 1.3 0.025 μm^2^, Nrxn3a and Nrxn3b RNA FISH particles: 0.03 μm^2^ and 0.01 μm^2^, zebrafish inner ear images – CTBP: 0.025 μm^2^, MAGUK: 0.025 μm^2^, mouse IHCs – CTBP: 0.025 μm^2^, GluR2: 0.025 μm^2^. A circularity factor between 0.1-0.5 was also applied to particle analysis. A watershed was applied to the particle analysis result to break apart overlapping synaptic components. After the watershed, the particle analysis was rerun with size and circularity thresholds to generate ROIs and measurements of each synaptic or RNA FISH component. The ROIs were applied to the original z-projection to get the average intensity and area of each punctum.

To recognize paired synaptic components, images were further processed using “Complete Synapse Counter v5.2”. Here, the overlap and proximity of ROIs from different channels (ex: pre- and post-synaptic puncta) was calculated. ROIs with positive overlap or ROIs within 2 pixels were counted as paired or partner components. The ROIs and synaptic component measurement (average intensity, area) and pairing results were then saved as Fiji ROIs, jpg images and csv files. For puncta counts the total number of ROIs was plotted per neuromast or per hair cell.

Some image datasets required a pre-processing step prior to entry into the “Complete Synapse Counter v5.2”. This includes zebrafish samples imaged at 3 dpf and our mouse IHCs datasets. For the pre-processing step, the volumes were segmented in VVDviewer (https://github.com/JaneliaSciComp/VVDViewer). Staining outside of the hair cell was manually segmented or removed using VVDviewer. After this segmentation, the z-stacks were then max-intensity projected and processed using the “Complete Synapse Counter v5.2” macro.

### Calcium imaging of lateral-line hair cells and afferents

For functional imaging, 4-6 dpf larvae were anesthetized in 0.04% Tricaine-S (tricaine methanesulfonate, Western chemical, TRS1), pinned to a Sylgard-filled perfusion chamber at the head and tail, and paralyzed by injection of 125 µM α-bungarotoxin (Tocris, 2133) into the heart cavity, as previously described (Lukasz and Kindt, 2018). Larvae were then rinsed three times in E3 embryo media to remove the tricaine. Next, larvae were rinsed three times with extracellular imaging solution (in mM: 140 NaCl, 2 KCl, 2 CaCl_2_, 1 MgCl_2_, and 10 HEPES, pH 7.3, OSM 310±10) and allowed to recover. Stimulation was achieved by a fluid jet, which consisted of a pressure clamp (HSPC-1, ALA Scientific) and glass pipette, pulled and broken to an inner diameter 40-50 µm, and filled with extracellular imaging solution. A 500-ms pulse of positive or negative pressure was used to deflect the hair bundles of mechanosensitive hair cells along the anterior-posterior axis of the fish. Hair cells of the two orientations (anterior and posterior) were stimulated separately. Stimuli that deflected kinocilia 5-15 µm were included in the analysis, as these deflections represent saturating stimuli that do not induce damage.

Hair-cell responses to stimuli were imaged with an A1R laser-scanning confocal scan head on an upright Nikon NI-E microscope with a resonant scanner and a 60x/1.0 NA CFI Fluor water immersion objective equipped with a z-piezo. Acquisition was controlled with Nikon Elements Advanced Research v. 5.20.02. GCaMP6s fluorescence was excited with a 488 nm solid-state laser passed through a standard 405/488/561/640 BS20/80 dichroic and collected with a 560 nm low-pass dichroic and 525/50 emission filter. Images were acquired using a GaAsP PMT and 4x averaging. Pixel size for presynaptic imaging was 0.28 µm; pixel size for MET imaging was 0.14 µm. Each neuromast (L2 or L3) was stimulated four times (starting with a posterior-to-anterior stimulus and alternating between the two directions) with an inter-stimulus interval of ∼2 min. This enabled us to collect presynaptic responses (collected first) and hair-bundle responses to both stimulus directions for each neuromast. 3 z-slices (1.5 µm step size for presynaptic responses; 0.5 µm step size for hair bundle responses) were collected per timepoint for 110 timepoints at a frame rate of 33 ms for a total of ∼100 ms per z-stack and a total acquisition time of ∼11 sec. Stimulation began at timepoint 31; timing of the stimulus was triggered by an outgoing voltage signal from Nikon Elements.

Calcium responses in the afferent process were acquired on a Swept-field confocal system built on a Nikon FN1 upright microscope (Bruker) with a 60x/1.0 NA CFI Fluor water-immersion objective. The microscope was equipped with a Rolera EM-C2 EMCCD camera (QImaging), controlled using Prairie view 5.4 (Bruker). GCaMP6s was excited using a 488 nm solid state laser. We used a dual band-pass 488/5611nm filter set (59904-ET, Chroma). Pixel size for postsynaptic imaging was 0.27 µm. Stimuli were delivered as outlined above for hair-cell responses. Each neuromast (L2, L3 or L4) was stimulated two times with an inter-stimulus interval of ∼2 min. 5 z-slices (1.0 µm step) were collected per timepoint for 80 timepoints at a frame rate of 20 ms for a total of ∼100 ms per Z-stack and a total acquisition time of ∼8 sec. Stimulation began at timepoint 31; timing of the stimulus was triggered by an outgoing voltage signal from Prairie view.

Acquired images were converted into TIFF series for processing. Researchers were blind to genotype during analysis. Z-stacks were average projected, registered, and spatially smoothed with a Gaussian filter (size = 3, sigma = 2) in custom-written MatLab software as described previously (Zhang et al., 2018). The first 10 timepoints (∼1 sec) were removed to reduce the effect of initial photobleaching. Registered average projections were then opened in Fiji for intensity measurements. Using the Time Series Analyzer V3 plugin, circular ROIs (18x18 pixels for presynaptic responses; 8x8 pixels for hair-bundle responses, 12x12 pixels for afferent process) were placed on hair bundles or synaptic sites; average intensity measurements over time were measured for each ROI, as described previously (Lukasz and Kindt, 2018). Neuromasts were excluded in the case of motion artifacts. Hair-bundles responses were excluded if they responded to stimuli of both directions. All other data was included in analyses. Presynaptic responses were defined as >10% ΔF/F0 within the 500 ms stimulus or >20% within 1 sec of stimulus onset. Hair-bundle responses were defined as >15% ΔF/F0 within the 500 ms stimulus and >15% in the 500 ms after the stimulus. Postsynaptic responses were defined as >5% ΔF/F0 and a minimum duration of 500 ms. Square wave responses indicate movement artifacts and were excluded. Calcium imaging data was further processed in Prism 10 (Graphpad). The first 20 timepoints were averaged to generate an F0 value, and all responses were calculated as ΔF/F0. Responses presented in figures represent average responses of synaptically active cells within a neuromast. The max ΔF/F0 was compared between wild-type animals and double mutants.

### Zebrafish startle behavior

A Zantiks MWP behavioral system was used to examine acoustic startle responses. Behavioral trials were performed at 5 dpf, on three independent days. For this behavioral analysis, we compared *nrxn3a^+/-^; nrxn3b^+/-^* double heterozygotes to *nrxn3a^-/-^; nrxn3b^-/-^* double mutants for an in-clutch, sibling comparison. *Nrxn3a^+/-^; nrxn3b^+/-^* double heterozygotes showed a slight (12%) yet significant reduction in complete synapses compared to wild-type controls. Therefore, we compared *nrxn3a^+/-^; nrxn3b^+/-^* double heterozygotes and *nrxn3a^-/-^; nrxn3b^-/-^* double mutants sibling to wild-type animals born the same day; this analysis revealed no difference in startle response between these genotypes.

The Zantiks system tracked and monitored behavioral responses via a built-in infrared camera at 30 frames per second. A 12-well plate was used to house larvae during behavioral analysis. Each well was filled with E3 and 1 larva. All fish were acclimated in the plate within the Zantiks chamber in the dark for 15 min before each test. To induce startle, an integrated stepper motor was used to drive a vibration-induced startle response. A vibrational stimulus that triggered a maximal % of animals startling in controls without any tracking artifacts (due to the vibration), was used for our strongest stimuli. Each larva was presented with each vibrational stimulus 5 times with 100 s between trials. For each animal, the proportion of startle responses out of the 5 trials was plotted. During the tracking and stimulation, a Cisco router connected to the Zantiks system was used to relay *x*, *y* coordinates of each larva every frame. To qualify as a startle response, a distance above 4 pixels or ∼1.9 mm was required within 2 frames after stimulus onset. Animals were excluded from our analysis if no tracking data was recorded for the animal.

### Experimental design and statistical analysis

Statistical analyses and data plots were performed with Prism 10 (Graphpad). Values of data with error bars on graphs and in text are expressed as mean ± SEM. A power analysis was performed to estimate approximate sample sizes needed. All zebrafish experiments were performed on a minimum of 4 animals, 7 neuromasts. Primary posterior lateral-line neuromasts with A-P orientations L1-L4 were used for all experiments except Ca_V_1.3 immunostains which examined L1, L2 and DV1 neuromasts. For 5 dpf larvae, each neuromast represents analysis from 12 to 20 hair cells and 41-68 synapses. For mouse studies all experiments were performed on at least 4 mutants and 4 siblings at P28 and P42. For synapse quantification, at least one ROI containing 8 IHCs were examined from each region of the cochlea (apex, mid, base) for each animal. All replicates are biological. Samples were scored and imaged blind to genotype whenever possible. Where appropriate, data was confirmed for normality using a D’Agostino-Pearson normality test. For pairwise comparisons, an unpaired t-test was used if data passed normality tests. If the data failed normality tests, a Mann-Whitney U test was used. For multiple comparisons, a one-way or two-way ANOVA was used.

### Code accessibility

The code Complete Synapse Counter v5.2 and Matlab software to visualize calcium signals will be deposited on Github.

---

**Fig S1.**
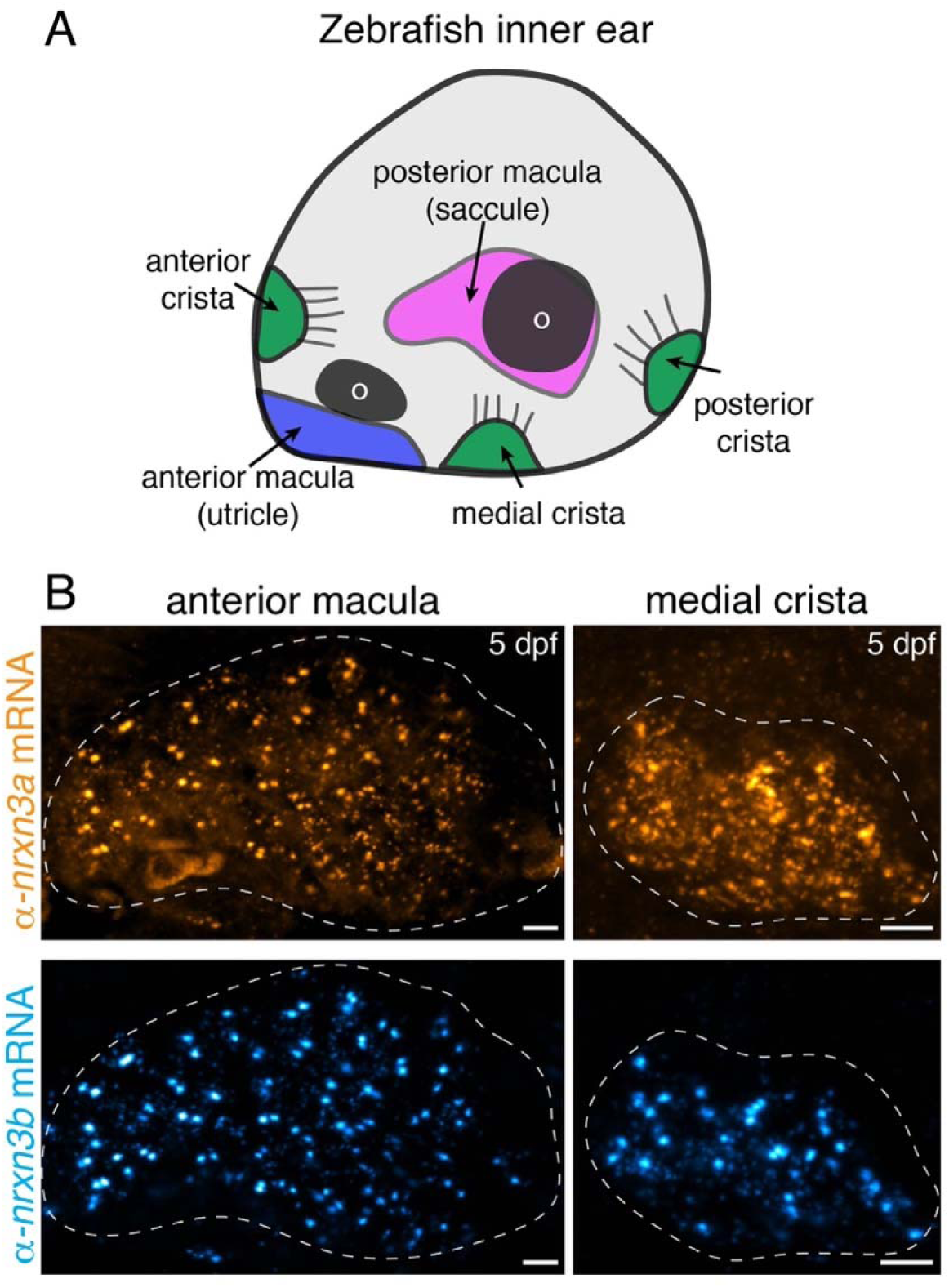
*nrxn3a* and *nrxn3b* mRNAs are present in zebrafish inner-ear hair cells. (**A**) Schematic showing a larval zebrafish inner ear. Within the inner ear, clusters of hair cells are present in 3 cristae and 2 maculae. Each macula is associated with an otolith (o). (**B**) RNA FISH analysis reveals that both *α-nrxn3a* (orange) and *α-nrxn3b* (cyan) mRNAs are present in inner-ear hair cells. The dashed line in B outlines the locations of hair cells within the sensory epithelium. Images are from larvae at 5 dpf. Scale bars = 5 µm in B.

**Fig S2.**
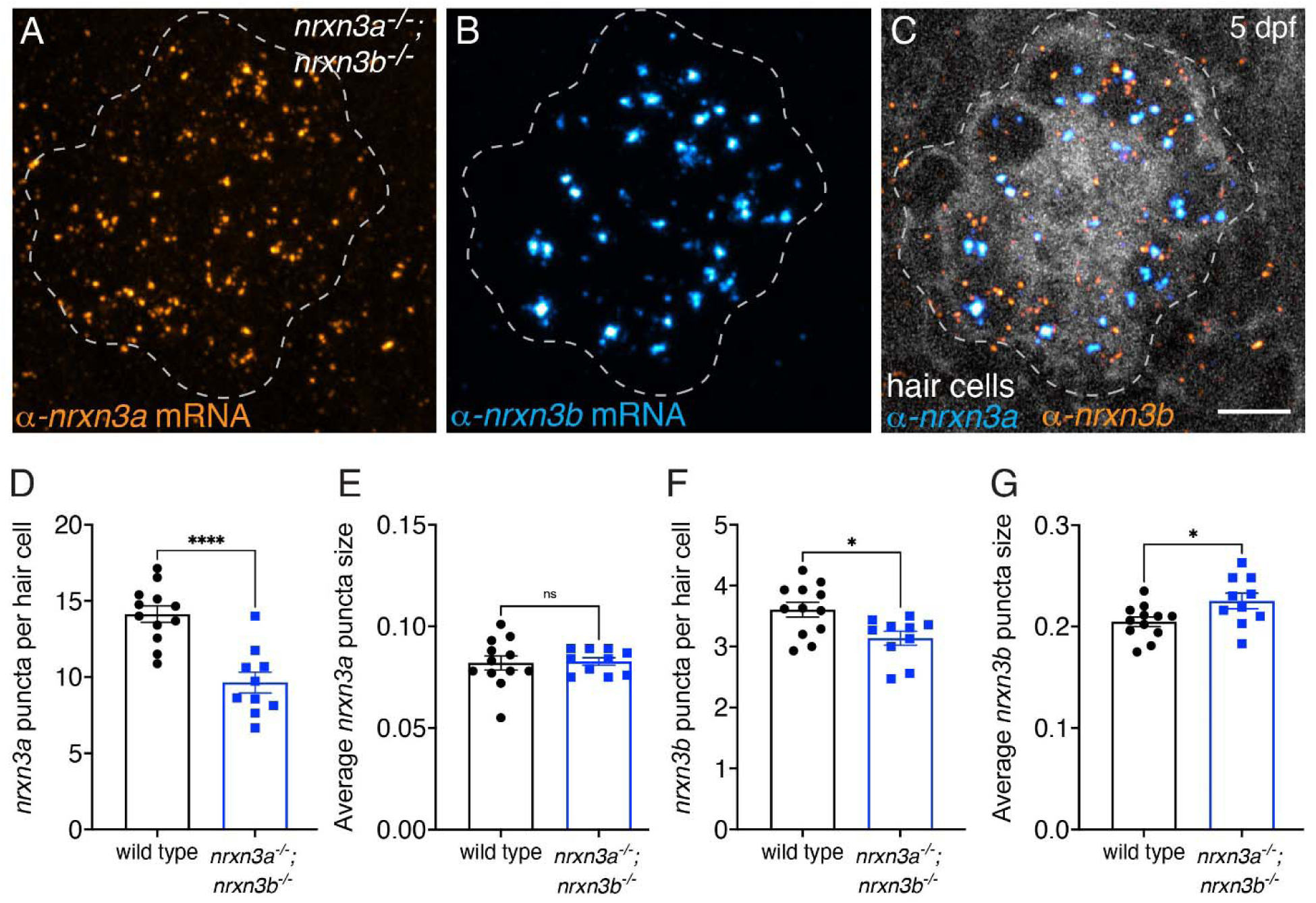
*nrxn3a* and *nrxn3b* mRNAs are reduced in lateral-line hair cells in zebrafish*nrxn3a; nrxn3b* mutants. (**A-C**) RNA FISH reveals that both *α-nrxn3a* (A, orange) and *α-nrxn3b* (B, cyan) mRNAs are present in lateral-line hair cells of *nrxn3a; nrxn3b* mutants. In C, hair cells (*myo6b:memGCaMP6s*) are labeled in grayscale. The dashed lines in A-C outline the locations of hair cells. (**D-G**) Quantification reveals that the number of *α-nrxn3a* (D) and *α-nrxn3b* (F) puncta are reduced in *nrxn3a; nrxn3b* mutants compared to wild-type controls. In addition, the size of *α-nrxn3b* (G), but not *α-nrxn3a* (E) puncta are slightly larger in *nrxn3a; nrxn3b* mutants compared to wild-type controls. An unpaired t-test was used in D-G, n = 12 wild-type and 10 *nrxn3a; nrxn3b* mutant neuromasts at 5 dpf. ns P > 0.05, *P < 0.05, ****P < 0.0001. Scale bar = 5 µm in C.

**Fig S3.**
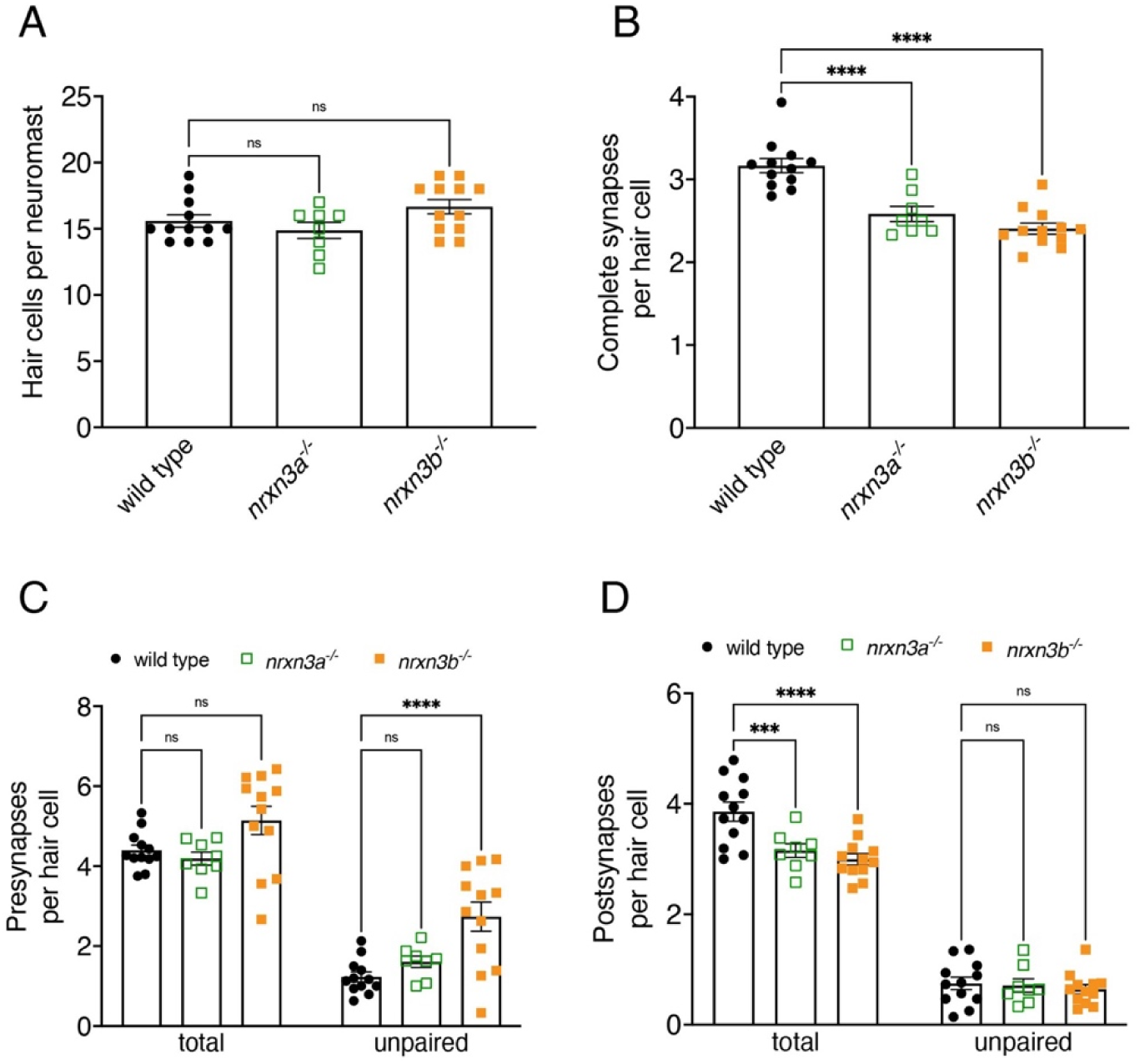
Minor defects in synapse organization are observed in *nrxn3a* and *nrxn3b* single mutants in mature hair cells at 5 dpf. (**A-F**) Quantification reveals that both *nrxn3a* and *nrxn3b* single mutants have a similar number of hair cells per neuromast compared to wild-type controls (A). There are significantly fewer complete synapses per hair cell in *nrxn3b* and *nrxn3a* single mutants compared to wild-type controls (B). The total number of pre-synapses are the same across all genotypes but there are significantly more unpaired presynapses in *nrxn3b* mutants (C). The total number of postsynapses per hair cell is significantly reduced in both in *nrxn3b* and *nrxn3a* single mutants compared to wild-type controls. In contrast, the number of unpaired postsynapses per hair cell is the same across all genotypes (D). N = 12 wild-type, 8 *nrxn3a* and 12 *nrxn3b* mutant neuromasts in A-D at 5 dpf. A one-way ANOVA was used in A-B, while a 2-way ANOVA was used in C-D. ns P > 0.05, *P < 0.05, ***P < 0.001, ****P < 0.0001.

**Fig S4.**
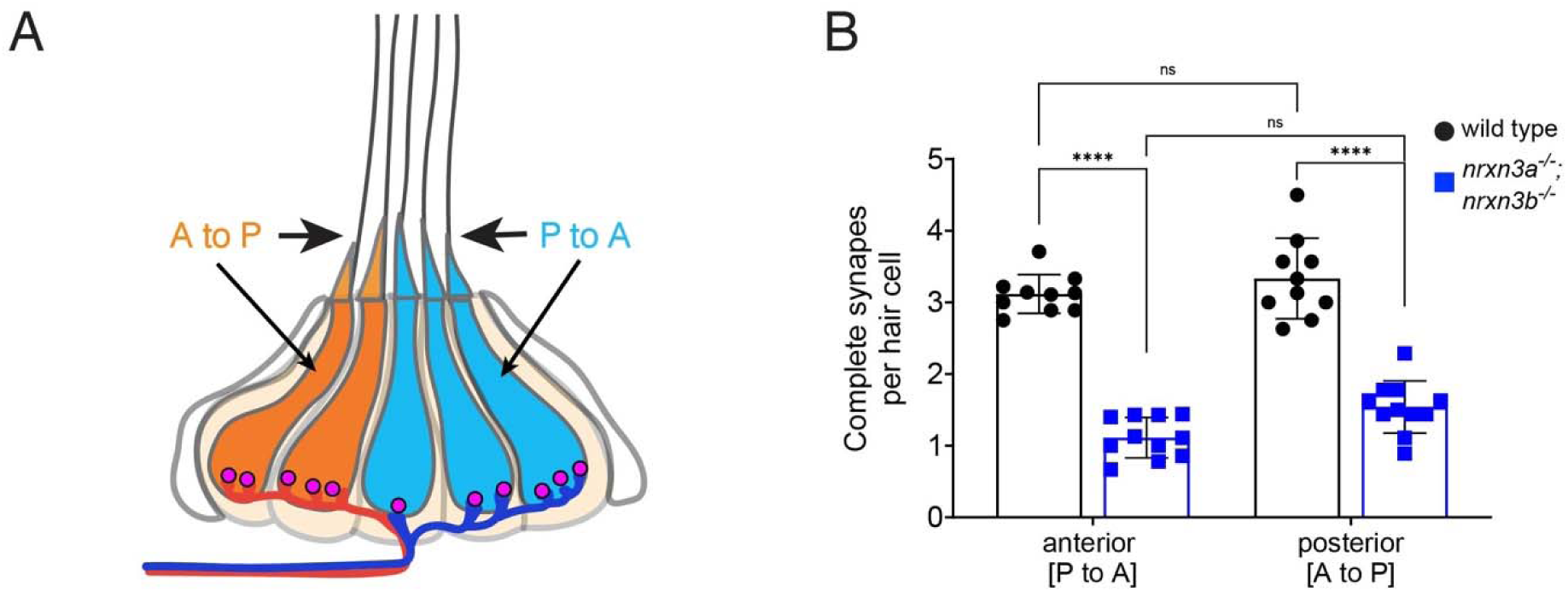
Synapse loss in mature hair cells is not linked to hair-cell orientation. **(A-B)** In primary posterior lateral-line neuromasts there are two populations of hair cells. One responds to anterior flow (blue, A), while the other responds to posterior flow (orange, A). Each population is selectively innervated by distinct afferent neurons (blue and orange processes). There is a significant and equivalent reduction in the number of complete synapses in hair cells that respond to anterior and posterior flow in *nrxn3a; nrxn3b* mutants compared to wild-type controls (B). N = 10 wild-type and 11 in *nrxn3a; nrxn3b* mutant neuromasts at 5 dpf. A 2-way ANOVA was used in B. ns P > 0.05, ****P < 0.0001.

**Fig S5.**
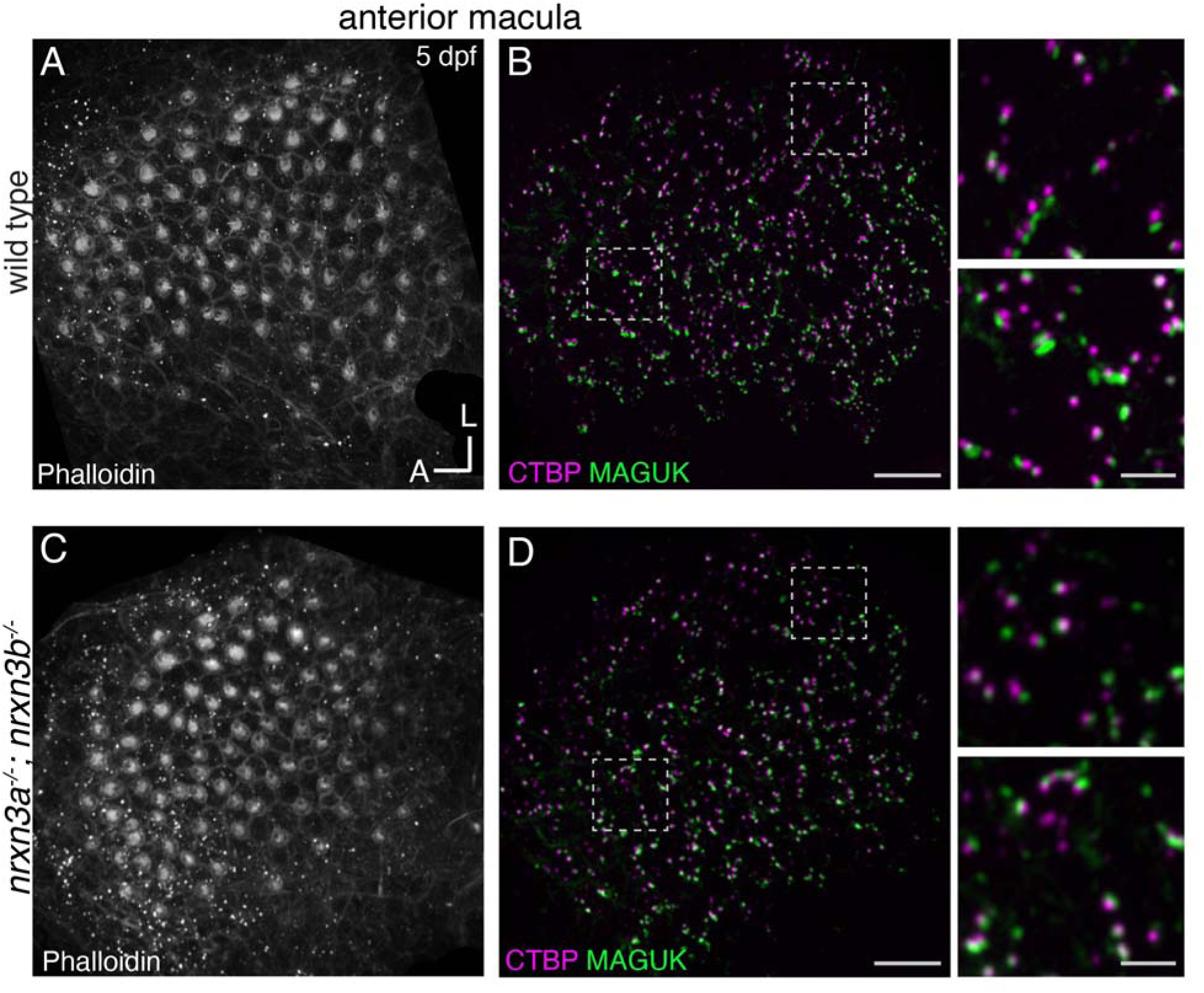
Loss of Nrxn3 results in fewer synapses hair cells in the zebrafish anterior macula. (**A-D**) Confocal images of the anterior macula (utricle) in wild-type controls (A-B, top panels) and *nrxn3a; nrxn3b* mutants (C-D, bottom panels). Phalloidin labels the apical hair bundles (A,C) while CTBP labels the presynapses (magenta), and MAGUK labels postsynapses (green) in B and D. The dashed lines indicate regions used to create the insets on the right side of panels B and D. The higher magnification insets show that there are fewer complete synapses in *nrxn3a; nrxn3b* mutants compared to wild-type controls. Images were taken from larvae at 5 dpf. Scale bars = 10 µm in A-D, 5 µm in the insets.

**Fig S6.**
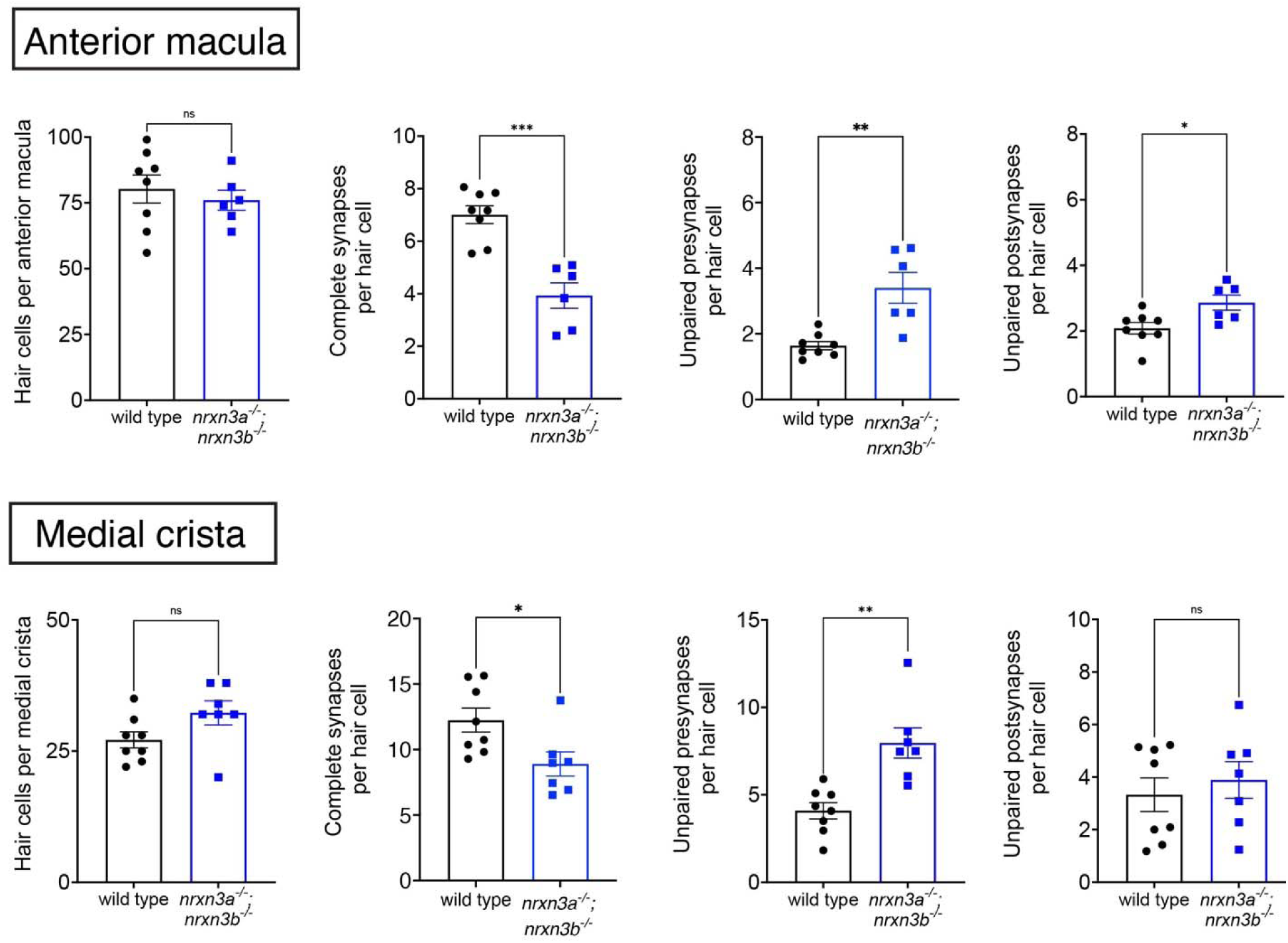
Quantification of synapse loss in anterior macula and medial crista in*nrxn3a; nrxn3b* mutants. Quantification reveals that wild-type controls and *nrxn3a; nrxn3b* mutants have a similar number of hair cells per anterior macula and medial crista. There are significantly fewer complete synapses per hair cell in each epithelium in *nrxn3a; nrxn3b* mutants compared to wild-type controls. Along with fewer complete synapses, there are significantly more unpaired presynapses per hair cell in *nrxn3a; nrxn3b* mutants compared to wild-type controls in both the anterior macula and medial crista. There are also more unpaired postsynapses per hair cell in the anterior macula, but not the medial crista in *nrxn3a; nrxn3b* mutants compared to wild-type controls. N = 8 wild-type and n = 6 *nrxn3a; nrxn3b* mutant anterior maculae, n = 8 wild-type and n = 7 *nrxn3a; nrxn3b* mutant medial cristae. Quantifications are from larvae at 5 dpf. An unpaired t-test was used for comparisons. ns P > 0.05, *P < 0.05, **P < 0.01, ***P < 0.001.

**Fig S7.**
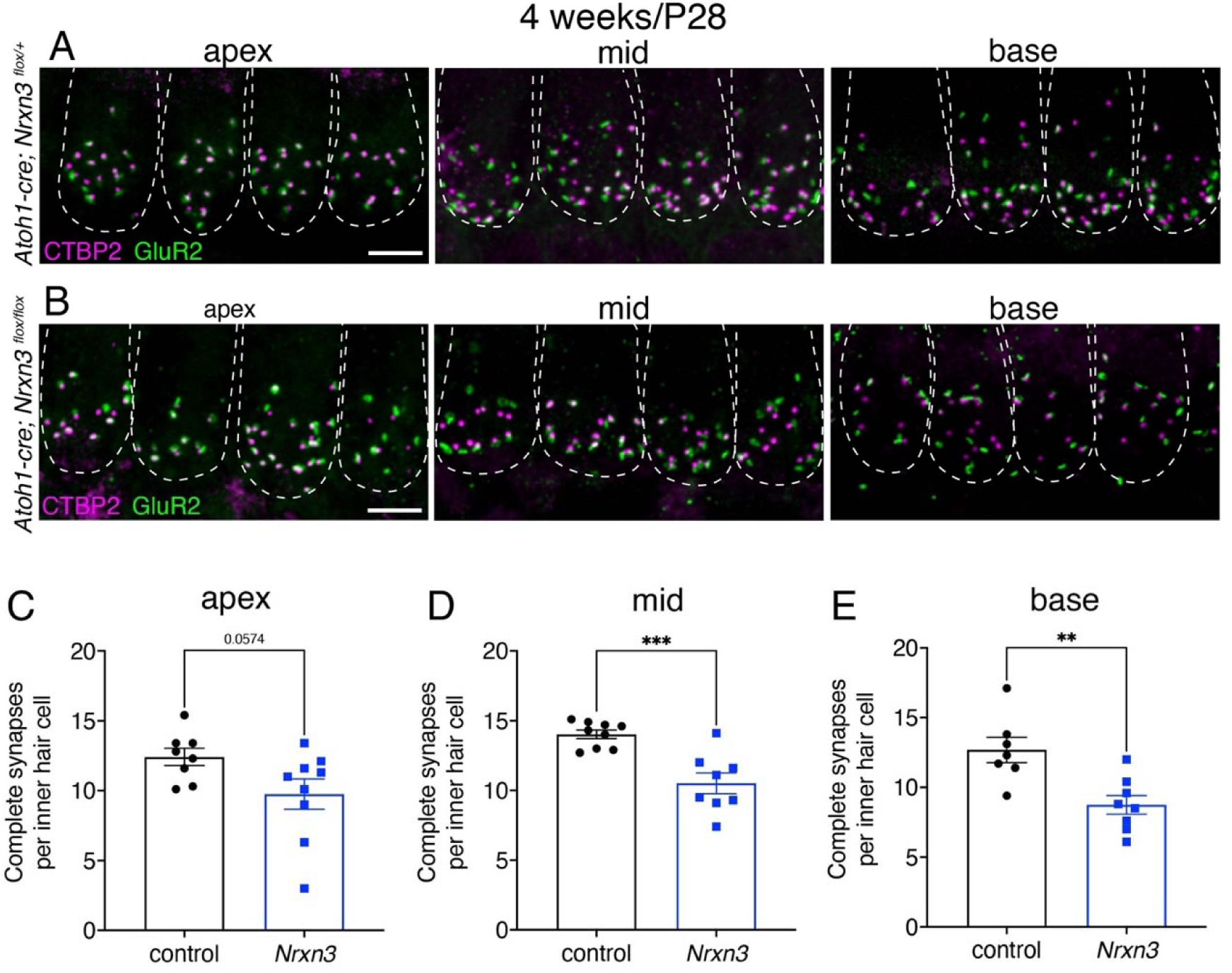
NRXN3 is required at 4 weeks for proper synapse numbers in mouse auditory inner hair cells. (**A-B**) Confocal images of 4-week old (P28) mouse inner hair cells from control (A) and *Nrxn3* mutant animals (*Atoh1-Cre; Nrxn3^flox/flox^*) (B). CTBP2 is used to label the presynapses (magenta), and GluR2 is used to label the postsynapses (green). Merged images show 4 IHCs from 3 different regions of the cochlea (apex, middle, basal thirds) for each genotype. Dashed lines indicate the outlines of hair-cell bodies in each image. (**C-E**) Quantification reveals that compared to controls, *Nrxn3* mutants have significantly fewer complete synapses per inner hair cell at the mid (D) and base (E), and a reduced but not significant decrease at the apex (C). N = 61 control and 58 *Nrxn3* IHCs for the apex region, 65 control and 59 *Nrxn3* IHCs for the for mid region, 50 control and 60 *Nrxn3* IHCs for the for base region. These findings were compiled from 4 animals from each genotype. An unpaired t-test was used in C-E. **P < 0.01, ***P < 0.001. Scale bar = 5 µm in A.

**Fig S8.**
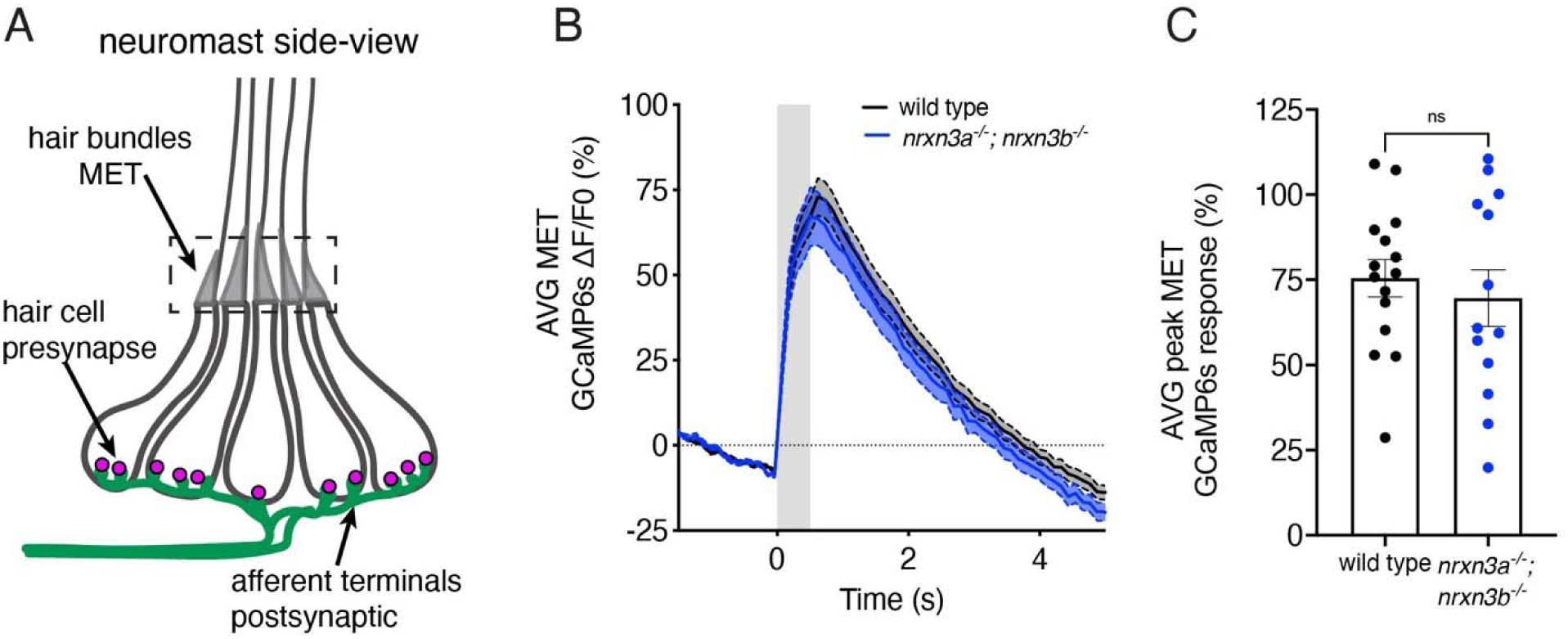
Loss of Nrxn3 does not impact the magnitude of mechanosensitive responses in lateral-line hair cells. **(A)** Schematic of a neuromast shown from the side. The region used to measure mechanosensitive GCaMP6 responses (MET) in apical hair bundles is indicated with a dashed box. (**B**) ΔF/F0 GCaMP6s traces showing average MET GCaMP6 response during a 500 ms fluid-jet stimulation (grey area) for wild-type controls (black) and *nrxn3a; nrxn3b* mutants (blue). Traces are displayed as mean, dashed lines are SEM. (**C**) Maximum ΔF/F0 MET calcium GCaMP6 during stimulation for wild-type controls (black) and *nrxn3a; nrxn3b* mutants (blue). N = 15 wild-type and 13 *nrxn3a; nrxn3b* mutant neuromasts at 5-6 dpf. An unpaired t-test was used in C. ns P > 0.05.

**Fig S9.**
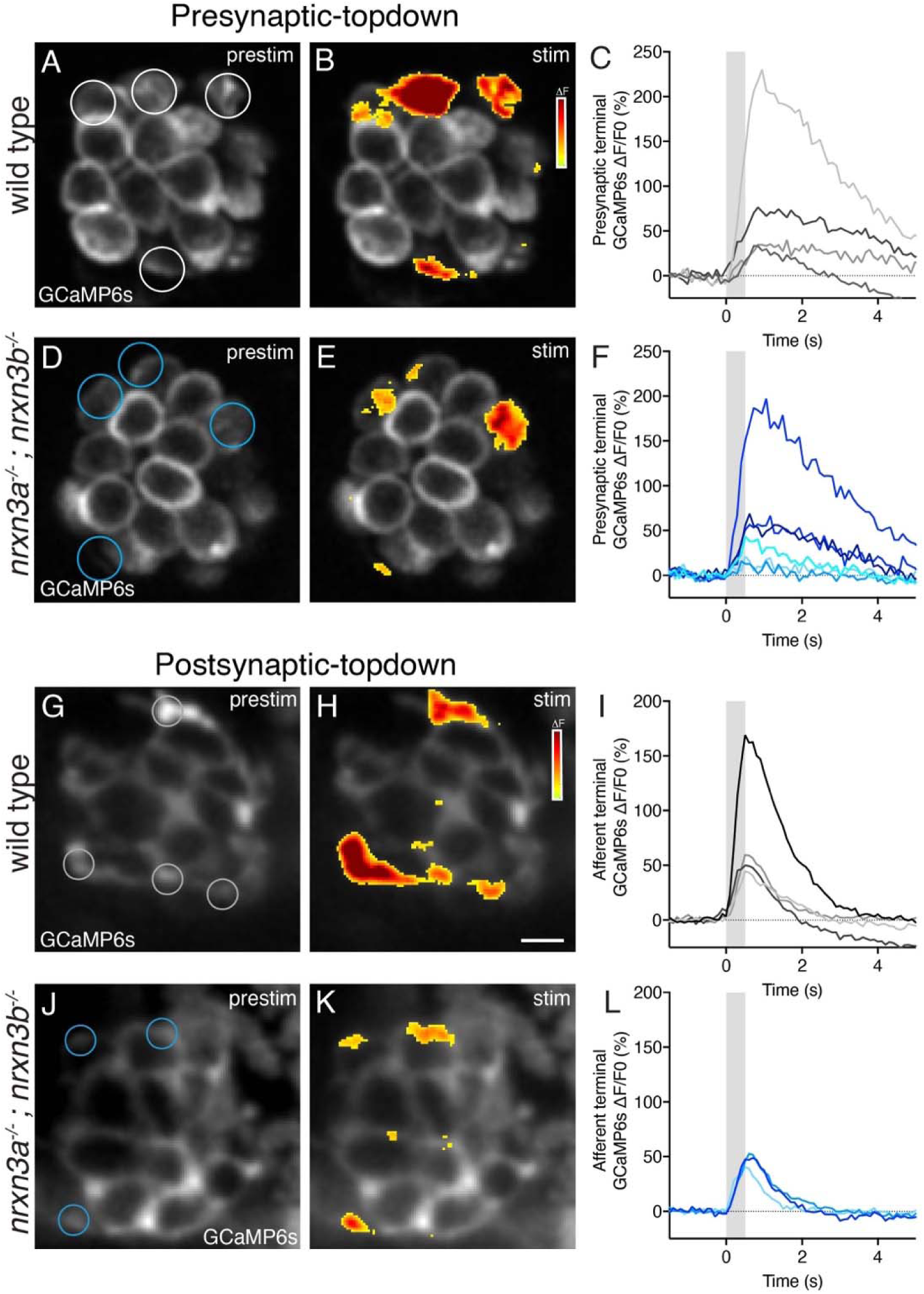
Nrxn3 is required for proper hair-cell synapse function in the lateral line. (**A-F**) ΔF heatmaps show spatial patterns of presynaptic GCaMP6s increases in hair cells before (A,D) and during (B,E) a 500 ms fluid-jet stimulation in a wild-type (A,B) and *nrxn3a; nrxn3b* mutant (D,E) neuromast. ROIs indicate synaptically active hair cells and examples of regions used to measure the average response per neuromast. Traces in C and F show ΔF/F responses from ROIs in A and D. Gray area indicates timing of stimulus. (**G-L**) ΔF heatmaps show spatial patterns of postsynaptic GCaMP6s increases in hair cells before (G,J) and during (H,K) a 500 ms fluid-jet stimulation in a wild-type (G,H) and *nrxn3a; nrxn3b* mutant (J,K) neuromast. ROIs indicate synaptically active terminals and examples of regions used to measure the average terminal response per neuromast. Traces in I and J show ΔF/F responses from ROIs in G and J. Gray area indicates timing of stimulus. Wild-type examples in A-B and G-H correspond to the same example in Fig 6 C,D and G,H.

